# Mapping the host protein interactome of non-coding regions in SARS-CoV-2 genome

**DOI:** 10.1101/2021.06.19.449092

**Authors:** Liuyiqi Jiang, Mu Xiao, Qing-Qing Liao, Luqian Zheng, Chunyan Li, Yuemei Liu, Bing Yang, Aiming Ren, Chao Jiang, Xin-Hua Feng

**Author notes:** Correspondence (M.X.), (L.Z.), (B.Y.), (A.R.), (C.J.), (X.F.). These authors contributed equally (Liuyiqi Jiang, Mu Xiao, Qing-Qing Liao, Luqian Zheng).

## Abstract

A deep understanding of SARS-CoV-2-host interactions is crucial to the development of effective therapeutics. The role of non-coding regions of viral RNA (ncrRNAs) has not been scrutinized. We developed a method using MS2 affinity purification coupled with liquid chromatography-mass spectrometry (MAMS) to systematically map the interactome of SARS-CoV-2 ncrRNA in different human cell lines. Integration of the results defined the core and cell-type-specific ncrRNA-host protein interactomes. The majority of ncrRNA-binding proteins were involved in RNA biogenesis, protein translation, viral infection, and stress response. The 5′ UTR interactome is enriched with proteins in the snRNP family and is a target for the regulation of viral replication and transcription. The 3′ UTR interactome is enriched with proteins involved in the cytoplasmic RNP granule (stress granule) and translation regulation. We show that the ORF10 is likely to be a part of 3′ UTR. Intriguingly, the interactions between negative-sense ncrRNAs and host proteins, such as translation initiation factors and antiviral factors, suggest a pathological role of negative-sense ncrRNAs. Moreover, the cell-type-specific interactions between ncrRNAs and mitochondria may explain the differences of cell lines in viral susceptibility. Our study unveils a comprehensive landscape of the functional SARS-CoV-2 ncrRNA-host protein interactome, providing a new perspective on virus-host interactions and the design of future therapeutics.

## Introduction

COVID-19 is an unprecedented global health threat caused by the severe acute respiratory syndrome coronavirus 2 (SARS-CoV-2)^1^. As of May 26, 2021, more than 160 million people have been infected, with more than 3.4 million deaths globally^2^. A detailed understanding of the molecular determinants of viral pathogenesis is urgently needed to provide more insights into the biology of SARS-CoV-2 and reveal potential therapeutic targets.

SARS-CoV-2 is an enveloped, positive-sense, and single-stranded RNA virus with a large genome of approximately 30 kb. The genomic structure of SARS-CoV-2 includes 14 open reading frames (ORFs). The largest ORF (ORF1a/b) encodes 16 nonstructural proteins (nsps) required for viral RNA productions^3^. The remaining ORFs encode nine accessory proteins and four structural proteins: spike (S), envelope (E), membrane (M), and nucleocapsid (N)^4,5^.

Consistent with known RNA viruses, SARS-CoV-2 relies on host proteins for assembling the replication and translation machinaries^6^. The genomic RNA (gRNA) serves as a dual-purpose template: 1. for the synthesis of the full-length negative-sense RNAs for genome replication and 2. for the synthesis of diverse subgenomic negative-sense RNAs (-sgRNAs) to make respective subgenomic mRNAs. During transcription, a set of 3′ and 5′ co-terminal-sgRNAs is generated by discontinuous transcription (Figure 1A)^7,8^. The discontinuous transcription involves a template switch from the body transcription regulatory sequence (TRS-B) to the leader TRS (TRS-L), located at about 70 nucleotides from the 5′ end of the genome (Figure 1A)^7,9^. To accomplish this, SARS-CoV-2 must employ unique strategies to utilize the host cell proteins while evading the host immune system. Thus, a thorough understanding of the interactions between viral RNAs and host proteins is essential. Several studies have attempted to explore the SARS-CoV-2 RNA-protein interactome^10-13^. Schmidt et al. identified physical associations between the viral RNAs and host proteins in infected human cells, revealing key pathways relevant to infection by using RNA antisense purification and mass spectrometry^10^. By integrating the ChIRP-MS data with genome-wide CRISPR screen data, Ryan A. Flynn et al. demonstrated a physical and functional connection between SARS-CoV-2 RNA and host mitochondria^11^. However, these studies did not systematically investigate the roles of non-coding regions of viral RNA (ncrRNAs) in viral infection.

**Figure 1.**
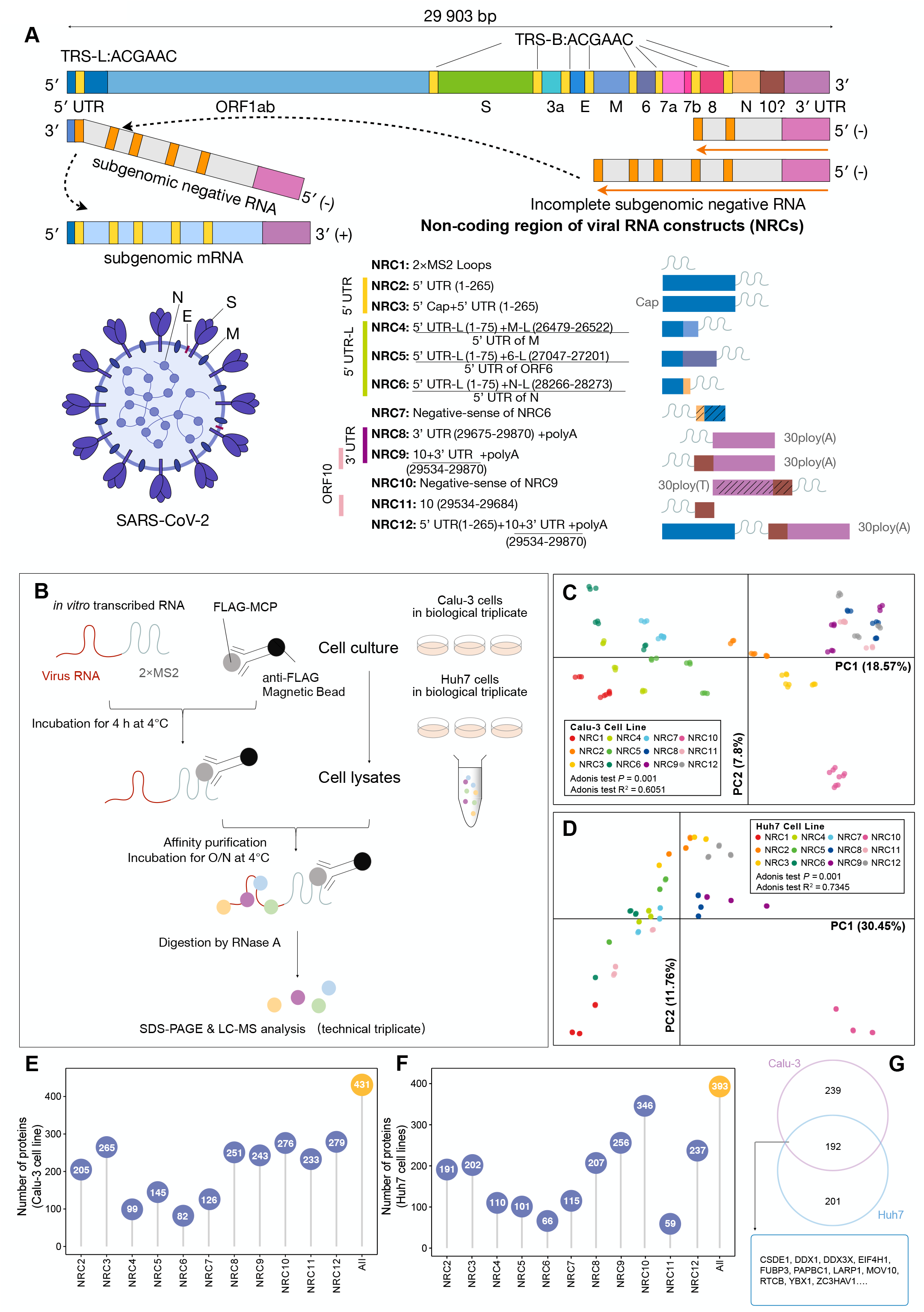
Mapping the SARS-CoV-2 ncrRNA interactome. (A) Schematic presentation of SARS-CoV-2 genome organization, the virion structure, discontinuous transcription, and ncrRNA constructs (NRCs). (B) The workflow of MS2 affinity purification coupled with liquid chromatography-mass spectrometry (MAMS) protocol. (C-D) PCA analyses showing the patterns of samples of NRCs in Calu-3 (C) and Huh7 (D). Dots represent samples. Colors indicate different NRCs. The adonis test was used to determine statistical significance. (E-F) Overview of the numbers of host proteins bound to different NRCs in Calu-3 (E) and Huh7 (F) cells. (G) Venn diagram of SARS-CoV-2 ncrRNA interactomes in Calu-3 and Huh7 cells.

In this study, we designed a series of MS2 linked viral ncrRNAs, utilized *in vitro* MS2 phage coat protein (MCP) affinity purification, and revealed the interacting host proteins through liquid chromatography-mass spectrometry. To identify the core and cell-specific interactome, we performed MAMS in human non-small-cell lung cancer cell line Calu-3 and hepatocellular carcinoma cell line Huh7, and further verified the interactions in a transduced human embryonic kidney-derived cell line HEK293T. In short, we identified an expanded SARS-CoV-2 ncrRNA interactome of 97 proteins and a core SARS-CoV-2 ncrRNA interactome of 52 proteins. We discovered that viral ncrRNAs had complex interactions with host proteins, including pro- and anti-viral factors. In particular, the 5′ UTR of viral genomic RNA was identified to bind to proteins involved in U1 snRNP and regulation of viral replication and transcription. The 3′ UTR had interactions with proteins involved in stress granule and translation regulation. We provide evidences that ORF10 is a part of 3′ UTR. We discovered that 5′ UTRs of subgenomic mRNAs may bind to different host proteins. Unexpectedly, the negative-sense ncrRNAs interacted with specific host proteins (e.g., CSTF3, RBM15, and SAMD9), indicating a potential role of negative-sense RNAs in infection. Finally, cell-type-specific interactome may provide a possible explanation for the variations in viral susceptibility of different cell lines.

## Results

### Developing MAMS to map the interactome between SARS-CoV-2 ncrRNAs and host proteins

We developed a method named MAMS (MS2 affinity purification coupled with liquid chromatography-mass spectrometry) to systematically map the interactome between SARS-CoV-2 ncrRNAs and host proteins. We constructed 12 ncrRNA constructs (NRCs) (Figure 1A). NRC1 comprises of MS2 and was designed as a global negative control. NRC2-NRC12 covers different types of ncrRNAs in SARS-CoV-2 (Figure 1A and Table S1). 5′ UTR and 3′ UTR are the two major non-coding regions of viral RNAs. Unlike eukaryotic cells, 5′ UTR and 3′ UTR of SARS-CoV-2 are ultra-conserved^14^. NRC2 and NRC3 were designed to investigate the interactome of the 5′ UTR; NRC8 and NRC9 were designed to investigate the interactome of the 3′ UTR. SARS-CoV-2 genome has nine TRS cores (5′-ACGAAC-3′), including one TRS-L and eight TRS-Bs, each TRS-B corresponds to an ORF (Figure S1A)^15,16^. Each subgenomic mRNA is joined between the TRS-L and the respective TRS-B (Figure 1A). The distance between TRS-Bs and respective ORFs showed significant differences, ranging from 0 to 155 bases (Figure S1A). We selected the TRS-Bs for N, M, and ORF6 as the others are too short. The three TRS-Bs were fused with the leader sequence of 5′ UTR (1-70 nt; referred to as 5′ UTR-L) to construct the complete 5′ UTRs of respective subgenomic mRNAs, namely NRC4, NRC5, and NRC6. The nature of ORF10 in the SARS-CoV-2 genome has been elusive and evidences show that ORF10 may not be a real coding region^16-19^. We therefore designed the NRC9 and NRC11 to explore the function of ORF10. To investigate the role of negative-sense ncrRNAs, we constructed the reverse-complementary of NRC9, namely NRC10, and the reverse-complementary of NRC6, namely NRC7. Finally, NRC12 comprising both 5′ UTR and 3′ UTR was synthesized to mimic the full length viral genomic RNA excluding the actual ORFs.

All NRCs were prepared by *in vitro* transcription at 37 °C with T7 RNA polymerase, followed by purification with denatured Urea-PAGE (polyacrylamide gel electrophoresis) and ethanol precipitation (Figure S1B). 3×FLAG-MCP was expressed as a recombinant protein in *Escherichia coli* BL21(DE3) Codon plus strain and purified with chromatography methods. The purified 3×FLAG-MCP was bound on anti-FLAG beads for MS2-tagged NRCs RNA binding (See “Methods” for details). The host proteins in cell lysates were incubated with respective NRCs to assemble NRC-protein complexes for affinity purifications. RNase A was then applied to release host proteins, which were then identified and quantified by liquid chromatography-mass spectrometry (LC-MS) (Figure 1B and S1C). We chose to map the interactions as the highly abundant viral proteins *in vivo* could easily mask interactions between low-abundance host proteins and viral ncrRNAs, thereby decreasing the sensitivity of detection. We performed the experiments in human lung cell line Calu-3 and human liver cell line Huh7, which are naturally susceptible to the SARS-CoV-2 but with different suceptibility^20^ (Figure 1B). All experiments were performed with three biological replicates and each biological replicate was quantified with three technical replicates to ensure the robustness of the approach (Figure 1B). We implemented data cleaning, data normalization^21^, and batch effect correction to address potential biases^22^ (Figure S2; see “Methods” for details). These considerations ensured rigorous and statistically meaningful analytical results.

### A comprehensive atlas of proteins bound to the SARS-CoV-2 ncrRNAs

Principal component analysis (PCA) and correlation analysis indicated that the results of experimental and technical replicates are highly consistent (Figure 1C-D and S3A-B; Spearman R = 0.90-0.99 and 0.74-0.99 in Calu-3 and Huh7 cells, respectively). The PCA clustering patterns reflect the design strategy in both cell lines. Briefly, samples of NRC2 and NRC3, both of which included 5′ UTR, were more similar; samples of NRC4, NRC5, and NRC6, all of which included 5′ UTR-L, were more similar; and samples of NRC8, NRC9, and NRC11, which included 3′ UTR or ORF10, were more similar (Figure 1C and D). These results indicated the high robustness and sensitivity of the approaches we used. Interestingly, we noticed a clear difference between samples of negative-sense NRC10 and the rest NRCs, while NRC7, the other negative-sense ncrRNA, did not show a similar separation.

To delineate the host proteins interacting with ncrRNAs of SARS-CoV-2, we first compared NRC2-NRC12 interactomes with the NRC1 interactome to identify statistically enriched proteins against the background (Wilcoxon test, FDR adjusted p-value < 0.05). A total of 431 (Calu-3) and 393 (Huh7) host proteins were identified to bind the SARS-CoV-2 ncrRNAs (Figure 1E and F; Table S2). 192 host proteins were identified in both cell lines (Figure 1G). These proteins may play essential roles in viral infection across cell lines. In addition, we compared the SARS-CoV-2 ncrRNA interactome to recently published SARS-CoV-2 RNA interactome in Huh7 cell line^10^. We found that 18 (37.5%) and 24 (50.0%) out of the 48 interacting proteins (excluding viral proteins) identified in the RNA-protein interactome overlapped with the ncrRNA-protein interactome in Calu-3 and Huh7 cells, respectively (Table S3), indicating that SARS-CoV-2 ncrRNAs contribute prominently to the total viral RNA-host protein interactome.

Protein domain enrichment analysis revealed that the SARS-CoV-2 ncrRNA interactome of both cell lines harbor abundant RNA-binding domains such as RNA recognition motif (RRM), DEAD/DEAH box helicase, KH domain, and LSM domain (Figure S3C and D). Of note, a recent study showed a depletion of the DEAD/DEAH box helicase domain in the SARS-CoV-2 RNA-host proteins interactome, most likely due to the lack of the 5′ UTR region in the experimental design^13^.

We next analyzed the biological functions of SARS-CoV-2 ncrRNA-binding proteins. Reactome pathway enrichment analysis^23^ revealed that the majority of ncrRNA-binding proteins were involved in cellular processes important for mRNA biogenesis and protein synthesis, including mRNA splicing, RNA transport, RNA decay, and translation (Figure 2A). Interestingly, proteins related to influenza infection and selenoamino acid metabolism were enriched in both cell lines (Figure 2A). Interferons (IFNs) play a vital role in the defense against coronaviruses^24-27^. It has been reported that IFN-λ3 stimulation can cause the upregulation of selenoamino acid metabolism^28^. These results indicate that the viral ncrRNAs may directly elicit host’s immune defense to the virus.

**Figure 2.**
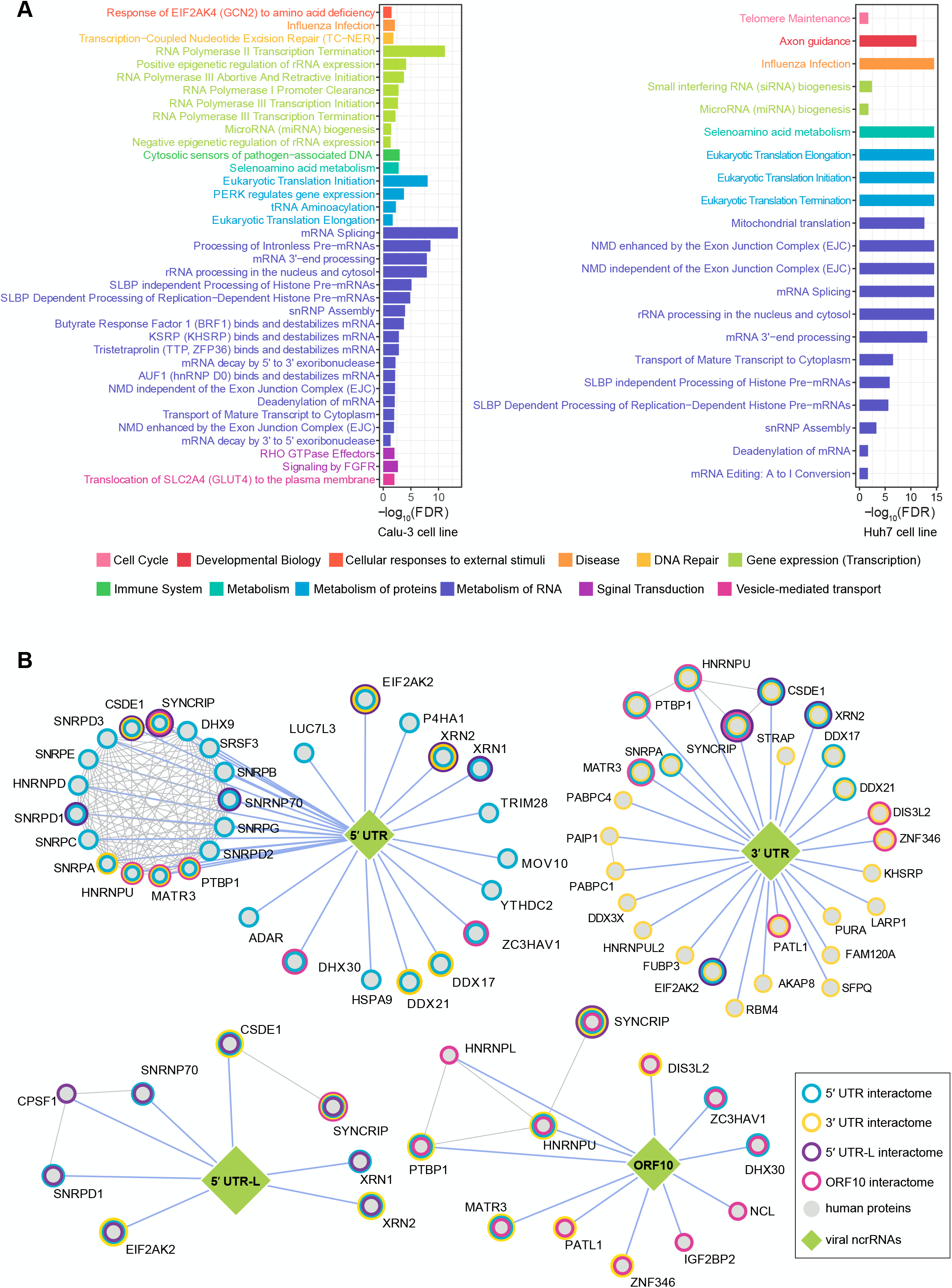
Analysis of the core SARS-CoV-2 ncrRNA interactome. (A) Reactome enrichment analysis demonstrating the major functions of the interactomes in Calu-3 (left) and Huh7 (right) cells. (B) Core interactome of SARS-CoV-2 ncrRNAs. Each node represents a host protein (closed circle) or viral ncrRNA (diamond). Color of nodes denotes the group of NRCs. Blue edges indicate the interactions between viral ncrRNAs and host proteins. Gray edges denote the interactions in host proteins.

### Expanded and Core SARS-CoV-2 ncrRNA interactomes

To dissect the biological functions of the viral ncrRNA interactome, we divided NRCs into four groups based on the design and PCA results (Figure 1A; 5′ UTR, 5′ UTR-L, 3′ UTR, and ORF10 groups). To find host proteins enriched in each group, we selected proteins consistently binding to NRCs in the same group across two cell lines (Wilcoxon test, FDR adjusted p-value < 0.05; Figure S3E and F). We visualized the interaction networks using the Cytoscape^29^ (Figure S4A). A total of 97 host proteins were identified as the expanded SARS-CoV-2 ncrRNA interactome (Figure S4A and Table S2). Notably, we observed that all proteins bound to 5′ UTR-L were enriched in the 5′ UTR group (Figure S4B), supporting the reliability of our approach. We then used a more stringent method to control within-group variation and compare samples of NRCs in the same group with the negative control NRC1 (Wilcoxon test, FDR adjusted p-value < 0.05, between-group variation/within-group variation > 2). As expected, the number of proteins identified across two cell lines by the stringent strategy decreased to 48 proteins (Figure S3E-F, S4C, and Table S2). To further validate these interactions, we performed RNA Binding Protein Immunoprecipitation (RIP) experiments in human kidney cell line HEK293T (Table S4; see “Methods” for details). The results of RIP experiments and mass-spec are highly consistent (Figure 2B and 3A). Integrating RIP data with mass-spec data, we finally identified 52 proteins as the core interactome (Figure 2B). In agreement with the expanded interactome, all proteins enriched in the core 5′ UTR-L interactome were identified to bind to the 5′ UTR (Figure 2B; CPSF1 was enriched in the expanded 5′ UTR interactome).

To further explore the functional importance, we intersected the core SARS-CoV-2 ncrRNA interactome with the published results of a genome-wide CRISPR screen designed to identify host proteins that impact SARS-CoV-2-induced cell death^11^. We obtained CRISPR data for 43 of the 52 proteins in the core ncrRNA interactome, including 26 proteins that significantly impact cell survival after SARS-CoV-2 infection (Figure 3B). Notably, SNRPC, AKAP8, and NCL had significant impacts on cell death. SNRPC and AKAP8 were involved in RNA splicing^30,31^. NCL may play a role in the process of transcription^32^. Two anti-viral factors, ZC3HAV1^33-35^and MOV10^36^, also impact on cell death (Figure 3B). ZC3HAV1 is known to inhibit the replication of different types of RNA viruses by degrading viral mRNAs, such as Ebola virus (EBOV) and Marburg virus (MARV)^33-35^. Other studies also reported that SARS-CoV-2 is restricted by ZC3HAV1^13,37^, reflecting a vital role of this protein during infection. MOV10 has been reported to bind to the SARS-CoV-2 RNA^10^. These results highlight the functional importance of proteins identified in the core ncrRNA interactome.

**Figure 3.**
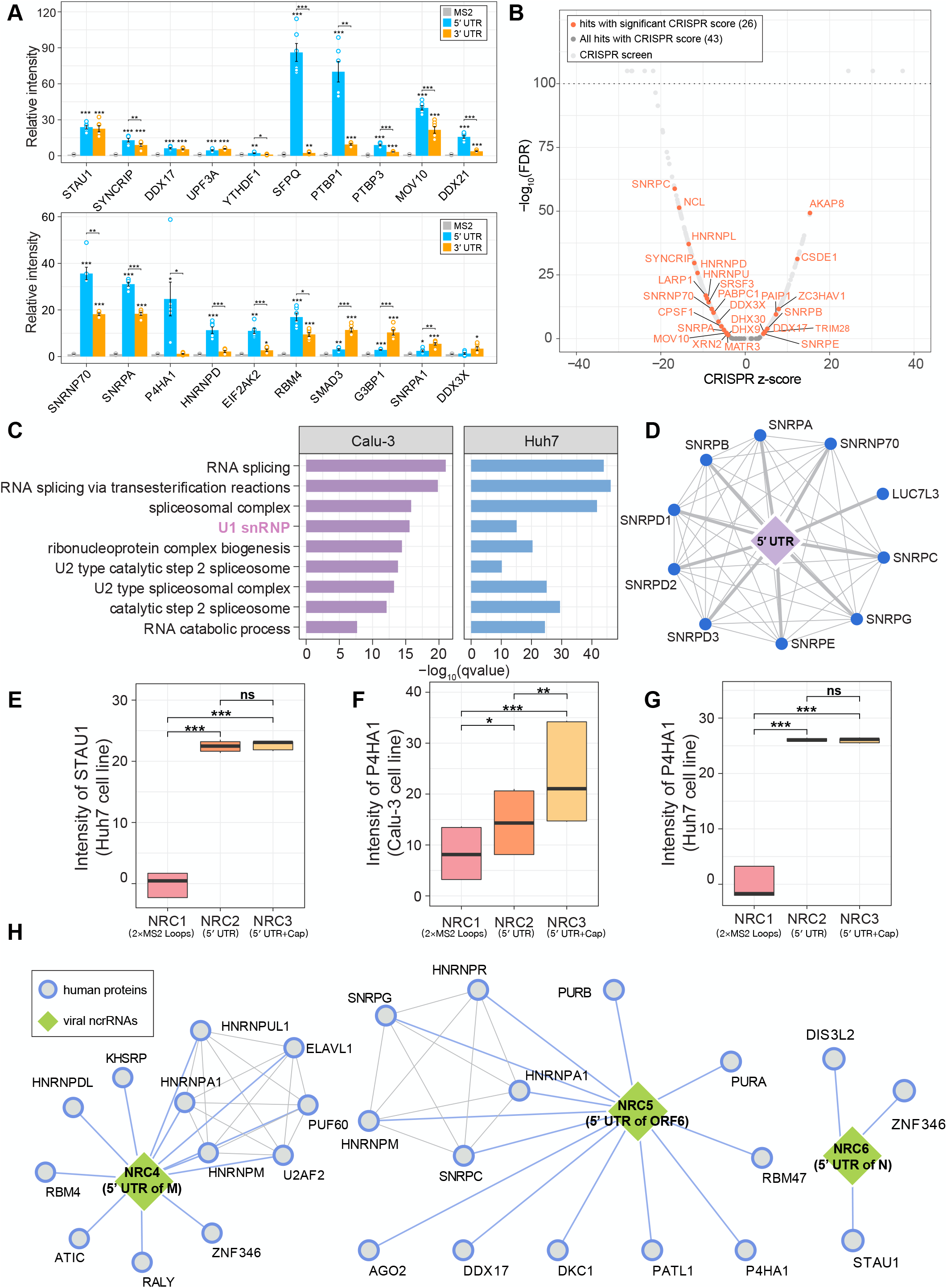
Functional characterization of the core SARS-CoV-2 ncrRNA interactome and 5′ UTR interactome. (A) RIP experiments validating the interactions between host proteins and SARS-CoV-2 5’ UTR or 3’ UTR. *P* values were calculated by the Wilcoxon test. * *p* < 0.05, ** *p* < 0.01, *** *p* < 0.001. (B) Volcano plot of interacting proteins in SARS-CoV-2 ncrRNA interactome overlaying with published CRISPR screen data in SARS-CoV-2-infected Vero E6 cells. Proteins with highly significant changes (-log10FDR >100) were not directly plotted. Orange color denotes proteins identified in SARS-CoV-2 ncrRNA interactome with significant changes (FDR < 0.05 and |CRISPR *z*-score| > 2). (C) GO enrichment results revealing the major functions of the 5**′** UTR interactome. (D) The interaction network of proteins involved in U1 snRNP. Each blue node represents a host protein. Edges represent the interactions between proteins. (E-G) The mass-spec intensity values of STAU1 (E) and P4H41 (F-G) in Calu-3 or Huh7 cells. *P* values were determined by Wilcoxon test. * *p* < 0.05, ** *p* < 0.01, *** *p* < 0.001. (H) Interaction networks of proteins interacted with the 5**′** UTRs of different subgenomic mRNAs. Each node represents a host protein (closed circle) or viral ncrRNA (diamond). Blue edges indicate the interactions between viral ncrRNAs and host proteins. Gray edges denote the interactions in host proteins.

### The 5′ UTR interacts with U1 snRNP and is a target for regulation of viral replication and transcription

To characterize the biological functions of the 5′ UTR interactome, we performed GO enrichment analysis which revealed that pathways associated with RNA splicing and RNA catabolic process were enriched in both cell lines (Figure 3C). RNA splicing pathways were also enriched in the 5′ UTR-L interactome (Figure S4D). In eukaryotes, U1, U2, U4, U6, and U5 small nuclear ribonucleoproteins (snRNPs) are components of the major spliceosome. However, the abundance of U1 snRNP in human cells far exceeds that of the other snRNPs^38^. Thus, we focused on the proteins involved in the U1 snRNP (Figure 3D). To confirm the interaction between U1 snRNP and 5′ UTR, we performed the RIP experiments using HEK293T cells. As shown in Figure 3A, SNRNP70 and SNRPA both had strong interactions with the 5′ UTR. These results supported that the 5′ UTR recruited proteins of U1 snRNP to assist the transcription and replication of SARS-CoV-2.

We discovered several proteins involved in the regulation of viral replication enriched in the 5′ UTR interactome. ADAR involved in SARS-CoV-2 genome editing^39^ interacted with the 5′ UTR (Figure 2B). We also noticed that STAU1, XRN1, and XRN2 had interactions with 5′ UTR (Figure 2B, 3A, and 3E). In RIP experiments, UPF3A was identified to bind to 5′ UTR. All four proteins are involved in RNA decay^40-42^, providing evidences that interactions between viral ncrRNAs and RNA decay proteins may be involved in host’s response to SARS-CoV-2 infection. SARS-CoV-2 attacks the lung and impairs gas exchange leading to systemic hypoxia, which activates the HIF-1 pathway^43^. P4HA1, which is reported to be essential for HIF-1α stabilization^44^, was enriched in the 5′ UTR interactome (Figure 3A, F, and G). These findings support that P4HA1 is a potential drug target for controlling lung damage elicited by viral infections. These results also highlight the necessity of further investigations into the hypoxia-related molecular mechanisms in SARS-CoV-2 infections.

Next, we compared the interactomes of 5′ UTR and 5′ UTR-L. RNA helicases are important for viral replication and transcription because both processes require RNA unwinding. Intriguingly, we found that 5′ UTR and 5′ UTR-L may bind to different sets of RNA helicases to accomplish the RNA unwinding process (Figure S4E). DDX5, DHX9, and DDX30 were identified to bind to 5′ UTR and 5′ UTR-L in both cell lines, demonstrating that the three RNA helicases may play important roles during the viral infection. However, in our data, the key enzyme involved in the formation of the 5′ cap structure, the guanylyltransferase, was not identified in the two NRCs without caps (NRC2 and NRC12). Altogether, these results indicate the important role of 5′ UTR in viral infection.

### Exploring the functional diversity of 5′ UTRs of subgenomic mRNAs

To explore potential similarities and differences in the 5′ UTRs of subgenomic mRNAs, we investigated proteins enriched in the interactomes of NRC4 (5′ UTR of ORF6), NRC5 (5′ UTR of M), or NRC6 (5′ UTR of N), but not shared among these constructs. We visualized the overlapping interactions in two cell lines (Calu-3 and Huh7) and discovered specific interactions between the NRCs and host proteins (Figure 3H). For example, our data revealed that ELAVL1 and RBM47 were enriched in the NRC4 and NRC5 interactomes, respectively. ELAVL1 (also named HuR) can regulate INF-β mRNA abundance and inhibit the type I IFN response^45^. RBM47 can promote the fish mitochondrial antiviral signaling protein degradation to suppress IFN production in zebrafish^46^.

We also used the RNAstructure^47^ to predict the secondary structure of NRC4, NRC5, and NRC6 (Figure S5A). Predicted results showed that the three NRCs shared the first two stem-loops (SL1 and SL2;1-60nt) conserved in all betacoronaviurs^48^, but had significant differences after 60 nt. These analyses suggest 5′ UTRs of different viral subgenomic mRNA could play different roles during transcription and translation.

### The 3′ UTR is involved in translation regulation and stress response

For SARS-CoV-2 and other coronaviruses, all viral mRNAs and genomic RNAs share the same 3′ UTR. Core proteins CSDE1, SYNCRIP, and EIF2AK2 were found in 3′ UTR interactomes (Figure 2B). CSDE1 plays an essential role in translational reprogramming, which regulates a number of RNAs^49^. Several studies demonstrate that CSDE1 can promote IRES-dependent translation initiation in viruses^50-52^. SYNCRIP may be directly involved in coronavirus mouse hepatitis virus (MHV) RNA synthesis as a positive regulator^53^. Thus, SARS-CoV-2 may recruit CSDE1 and SYNCRIP to promote viral replication and translation. In contrast, interferon-induced, double-stranded RNA-activated protein kinase EIF2AK2 can exert antiviral activity on a wide range of DNA and RNA viruses, including hepatitis C virus (HCV), hepatitis B virus (HBV), measles virus (MV), and herpes simplex virus 1 (HHV-1)^54-60^. The interactions between EIF2AK2 and viral ncrRNAs indicate that the EIF2AK2 represents the early immune response to viral infection and could be a therapeutic target. Notably, the three core proteins were also enriched in the 5’ UTR and 5′ UTR-L interactomes (Figure 2B), providing evidence that SARS-CoV-2 may use the “closed loop model” to control mRNA translation^61^. LARP1, FUBP3, and several DEAD-box RNA helicases were enriched in the 3′ UTR group (Figure 2B). A recent study reported that LARP1 can bind to the 3′ UTR of SARS-CoV-2^10^. FUBP3 can regulate mRNA localization by binding to the 3′ UTR of cellular mRNAs^62^. These results indicate the important role of 3′ UTR of SARS-CoV-2 during translation.

In agreement with the 5′ UTR interactome, GO enrichment analysis showed that the 3′ UTR interactome is enriched with pathways associated with RNA splicing and RNA catabolic process (Figure 4A). Notably, we discovered that the cytoplasmic stress granule (SG) GO term was enriched in both cell lines (Figure 4A and B). SGs can exert anti-viral functions via various mechanisms, including sequestrating host and viral mRNAs and proteins, recruiting immune signaling intermediates, and inhibiting protein synthesis^63-65^. G3BP1 is crucial to the formation of SGs^66^, although our mass-spec data did not find interactions between G3BP1 and the 3′ UTR, probably due to technical limitations, our RIP experiments confirmed that G3BP1 binds to 3′ UTR in HEK293 cells (Figure 3A). These observations demonstrate that host cells may try to block SARS-CoV-2 production via the interaction between SGs and 3′ UTR. Moreover, we found that Smad3 had a strong interaction with 3′ UTR in HEK293T and Calu-3 cells (Figure 3A and 4C). Smad3 plays an essential role in regulating the TGF-β pathway^67^. A previous study reveals that the N protein of SARS-CoV can associate with Smad3 to interfere with the complex formation between Smad3 and Smad4, blocking apoptosis of SARS-CoV infected host cells^68^. These results further support that TGF-β mediated responses can be a valid target for the treatment of COVID-19.

**Figure 4.**
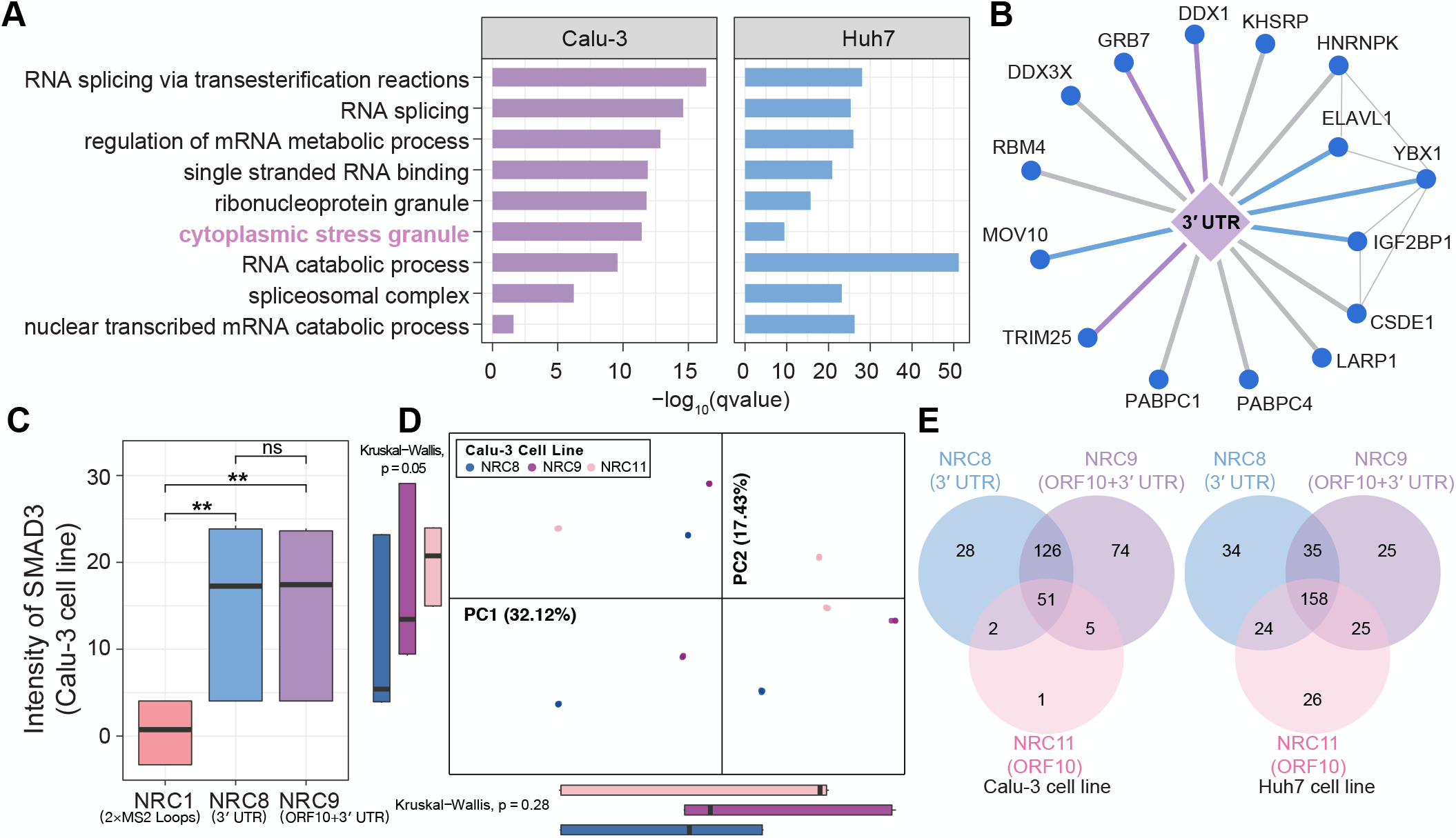
Functional characterization of interactome of the 3′ UTR. (A) GO enrichment results revealing the functions of the 3**′** UTR interactome. (B) The interaction network of proteins involved in the cytoplasmic stress granule. Each blue node represents a protein. Blue and purple edges denote interactions with proteins enriched in the 3**′** UTR interactome of Huh7 or Calu-3 cells, respectively. Gray edges denote interactions with proteins enriched in both cell lines. (C) The mass-spec intensity value of Smad3 in Calu-3 cells. *P* values were calculated by the Wilcoxon test. * *p* < 0.05, ** *p* < 0.01, *** *p* < 0.001. (D) PCA analysis of interactomes of the NRC8, NRC9, and NRC11 in Calu-3 cells. Each dot represents a sample. Colors represent different NRCs. *P* values were calculated by the Kruskal-Wallis test. (E) Venn diagram of the interactomes of NRC8, NRC9 and NRC11.

### ORF10 is likely a part of the 3′ UTR

To investigate the role of ORF10, we compared the interactomes of NRC8 (3′ UTR), NRC9 (ORF10 + 3′ UTR), and NRC11 (ORF10). PCA analysis showed no significant differences among the three NRCs in Calu-3 cells (Figure 4D). In Huh7 cells, PC1 revealed a significant separation among the three interactomes (Figure S5B). However, the interactome of NRC11 was more similar to the interactome of NRC8 instead of NRC9. Moreover, the interactomes of the three NRCs shared most of the proteins (Figure 4E). We also applied the GO enrichment analysis to the ORF10 interactome. The major functions of proteins bound to ORF10 included RNA splicing, mRNA metabolic process, and RNA catabolic process, showing high consistency with the 3′ UTR interactome (Figure S5C). Importantly, we found that most proteins bound to ORF10 were also enriched in the 3′ UTR group (Figure 2B). Of note, we observed that insulin-like growth factor 2 mRNA-binding protein (IGF2BP2) interacted with ORF10 (Figure 2B). IGF2BP2 is reported to play a key role in mRNA localization, stability, and translation control^69,70^. These results indicate that at least function-wise, ORF10 is very likely to be a part of the 3′ UTR.

### The emerging roles of negative-sense ncrRNAs during viral infections

Next, we asked if negative-sense ncrRNAs could play a role during infection. To this end, we first compared the interactomes of negative-sense ncrRNAs to the corresponding ncrRNAs. We found that most proteins bound to NRC6 or NRC9 were also enriched in NRC7 or NRC10 (negative-sense) interactomes (Figure 5A), but NRC7 and NRC10 bind to more proteins overall. Based on the PCA analysis of Calu-3 and Huh7 cells (Figure 1C and D), we focused on the interactomes of NRC10. Overall, specific interactions between NRC10 and host proteins were observed in both cell lines (Figure 5B). Several cleavage factors (CSTF3, CSTF2, and CPSF7) were found to specifically bind to NRC10. Remarkably, the m^6^A writer RBM15/RBM15B was identified to interact with NRC10 in Huh7 cells (Figure 5B). A recent study demonstrates that the m^6^A is prevalent in the negative-sense RNA of SARS-CoV-2^71^. In Calu-3 cells, SAMD9 was identified to interact with NRC10 (Figure 5B). Other studies showed that SAMD9 is an innate antiviral host factor and several viruses antagonize SAMD9 to prevent granule formation^72,73^.

**Figure 5.**
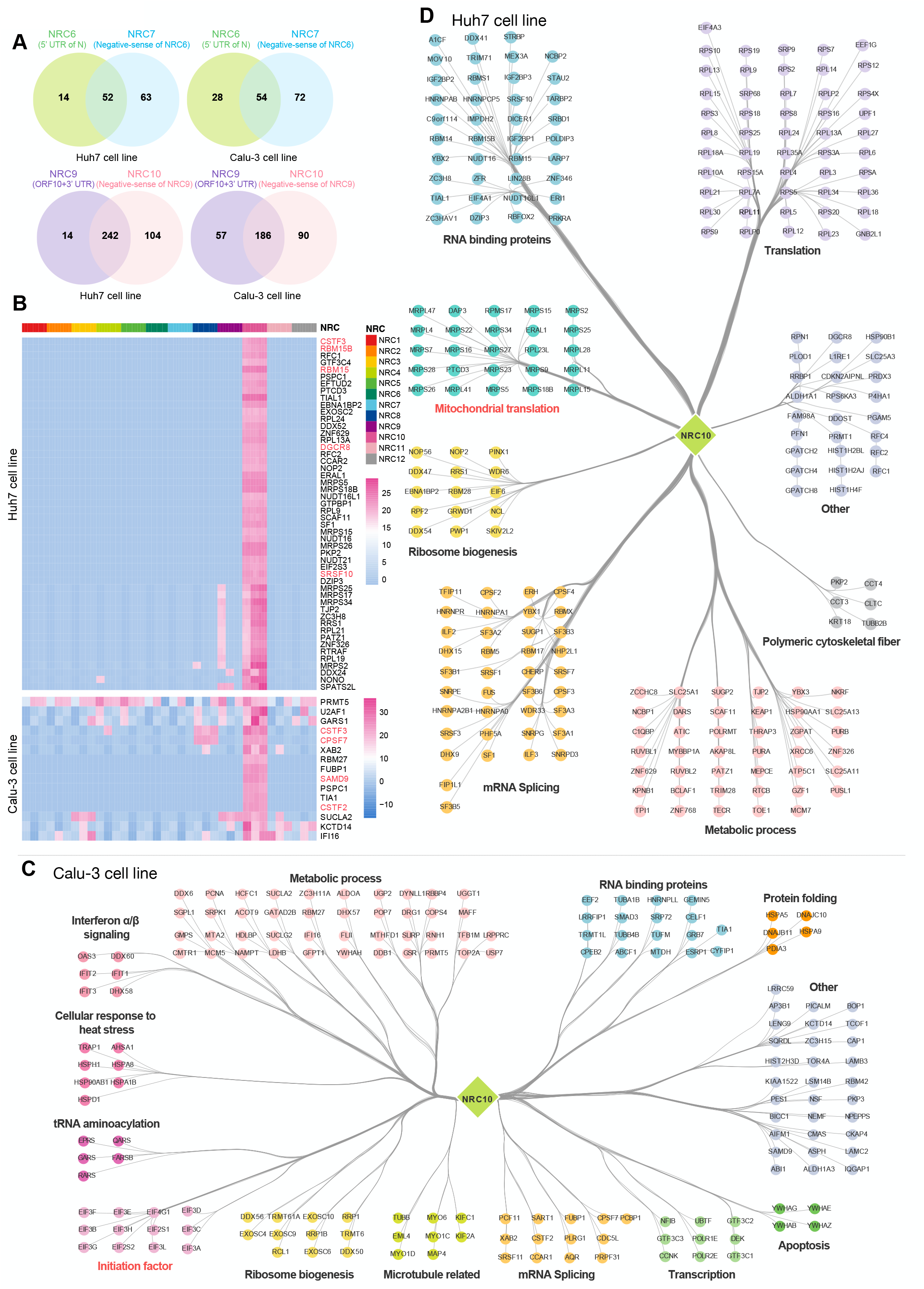
Interactome of the SARS-CoV-2 negative-sense ncrRNAs. (A) Venn diagram of interactomes of negative-sense ncrRNAs and corresponding ncrRNAs. (B) Heatmap showing proteins interacted with NRC10. (C-D) Protein interaction network of NRC10 in Calu-3 (C) and Huh7 (D) cells. Color indicates the category of biological functions.

We next constructed the interaction network of proteins bound to NRC10 in Calu-3 and Huh7 cells, respectively (Figure 5C and 5D), demonstrating a significant difference between the two cell lines. The NRC10 interactome in Calu-3 cells revealed an enrichment of proteins in the eukaryotic translation initiation factor 3 (eIF3) family (Figure 5C and S6A). Whilst in Huh7 cells, proteins involved in mitochondrial translation and peptide chain elongation were enriched in the NRC10 interactome (Figure 5D). These results opened new avenues to explore unknown regulatory mechanisms of viral negative-sense ncrRNAs in viral infection.

### Cell-type-specific patterns of SARS-CoV-2 ncrRNA-host protein interactome

SARS-CoV-2 infection leads to damages and dysfunctions in different types of organs and tissues, including damages to lung and liver^74,75^. Our previous study revealed that there are significantly different infection outcomes among cell lines^20^. Specifically, the human lung cell line Calu-3 is more susceptible to SARS-CoV-2 infection than the human liver cancer cell line Huh7. Therefore, we explored whether the cell-type-specific interactions could explain this phenotype. We discovered that 239 (55.45%) and 201 (51.15%) proteins were exclusively enriched in the interactomes of the Calu-3 and Huh7 cell lines, respectively (Figure 1G). In the four groups mentioned previously, the two cell lines exhibited a significant difference (Figure S6B). In addition, Reactome enrichment analysis revealed that the interactome of Calu-3 cells was associated with signal transduction (e.g., RHO GTPase effectors) and vesicle-mediated transport (Figure 2A). Increasing evidence shows that RHO GTPase signaling interferes with multiple steps in the viral replication cycle, and many viruses manipulate this signal transduction for their benefits^76^. Moreover, proteins involved in the PERK pathway were also enriched in the interactome of the Calu-3. A previous study reported that the Seneca Valley virus (SVV), a nonenveloped, single-stranded, and positive-sense RNA, could induce autophagy through the PERK pathway^77^.

GO enrichment analysis revealed that two cell lines had clear differences (Figure 6A). For example, the SARS-CoV-2 ncrRNA interactome of Huh7 displayed an increased enrichment of ribosome and nuclear-transcribed mRNA catabolic process nonsense-mediated decay (Figure 6A). Furthermore, we also compared the biological functions of interactomes of 5′ UTR, 5′ UTR-L, and 3′ UTR in the two cell lines. Notably, the GO terms related to nonsense-mediated decay was enriched in the interactome of 3′ UTR in Huh7 cells (Figure 6B); the apoptotic signaling pathway GO term were enriched in the interactome of 5′ UTR-L and 5′ UTR in Calu-3 cells (Figure 6C and S6C). Thus, we hypothesized that these pathways might be associated with the high infection ratio and cell death ratio of Calu-3 cells. Proteins detected in both cell lines also showed different patterns (Figure 6D). For example, PTBP3 and FAM120A bound to most NRCs in Calu-3 cells. However, in Huh7 cells, the two proteins only had interactions with selected few NRCs. Intriguingly, we observed an enrichment of mitochondrial ribosomal proteins in Huh7 cells but not in Calu-3 cells (Figure 6E). Altogether, these results reveal cell-type-specific interactions between SARS-CoV-2 ncrRNAs and host proteins, especially mitochondria proteins, suggesting mitochondria could play a crucial role during infection.

**Figure 6.**
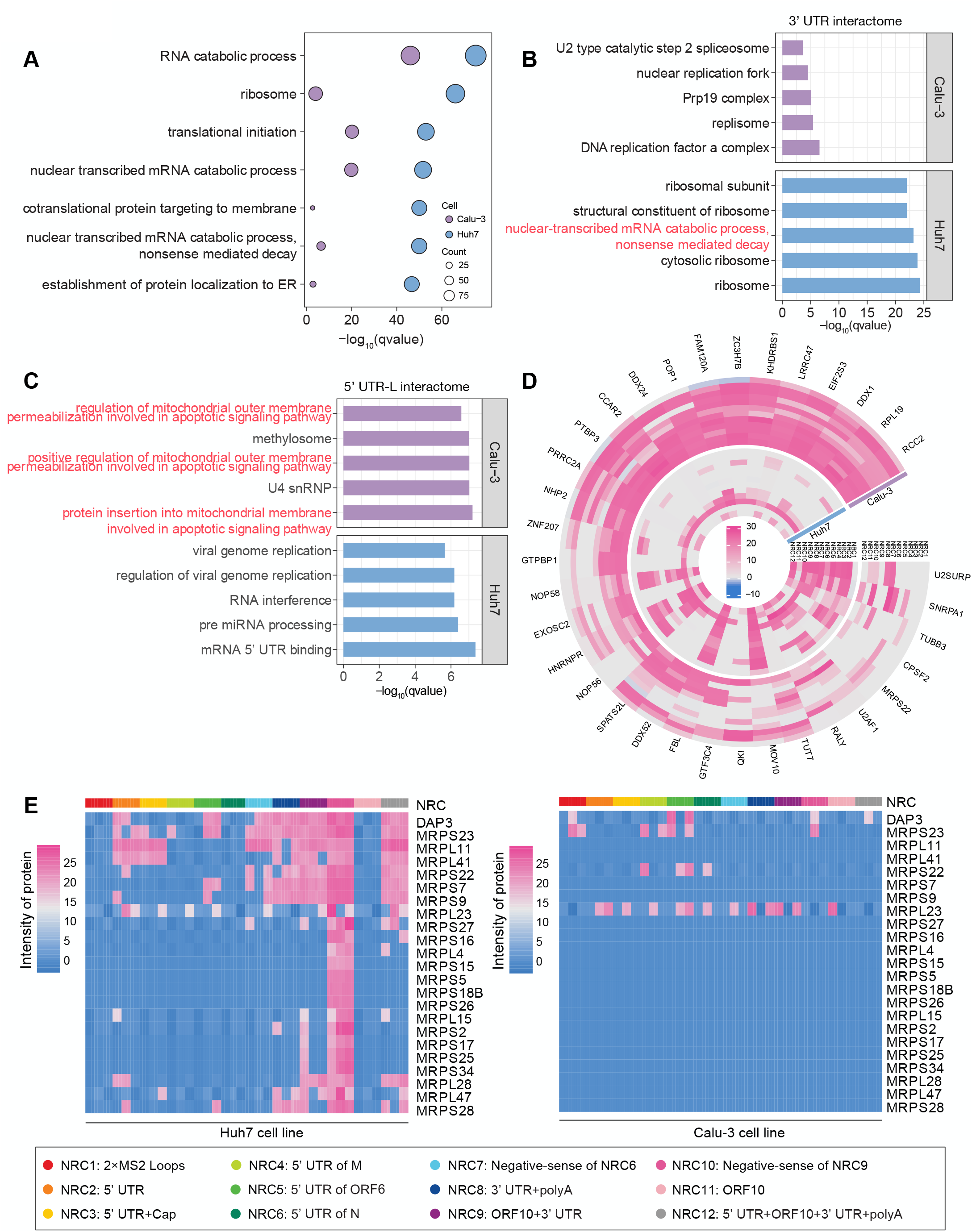
Cell-type-specific interactions between SARS-CoV-2 ncrRNA and host proteins. (A) GO enrichment revealing that SARS-CoV-2 ncrRNA interactomes are different in Calu-3 and Huh7 cells. (B-C) GO enrichment revealing different biological functions of the 3**′** UTR (B) and 5**′** UTR-L (C) interactomes. (D) Circular heatmap showing the cell-type-specific interactions between SARS-CoV-2 ncrRNAs and host proteins. (E) Heatmap showing that the mitochondrial ribosome proteins were specifically enriched for the SARS-CoV-2 ncrRNA interactome in Huh7 cells.

## Discussion

Decoding how the ncrRNAs of SARS-CoV-2 interact with host proteins is a crucial piece to the understanding of virus biology and COVID19. The present study provides a comprehensive view of the landscape of SARS-CoV-2 ncrRNA-host protein interactome (Figure 7). In this study, we developed MAMS to systematically map the viral ncrRNA-host protein interactome across cell lines. Comparing with previous studies, the MAMS approach effectively removed the masking effects of the high abundance of viral proteins during *in vivo* infection, and revealed significant more potential host proteins interacting with the viral ncrRNAs (Figure 1E-F and 2B), many of which were also validated by RIP experiments. Integration of the MAMS data from the Calu-3 and Huh7 cell lines and RIP data from the HEK293T cell line, we identified the core and cell-type-specific SARS-CoV-2 ncrRNA interactomes.

**Figure 7.**
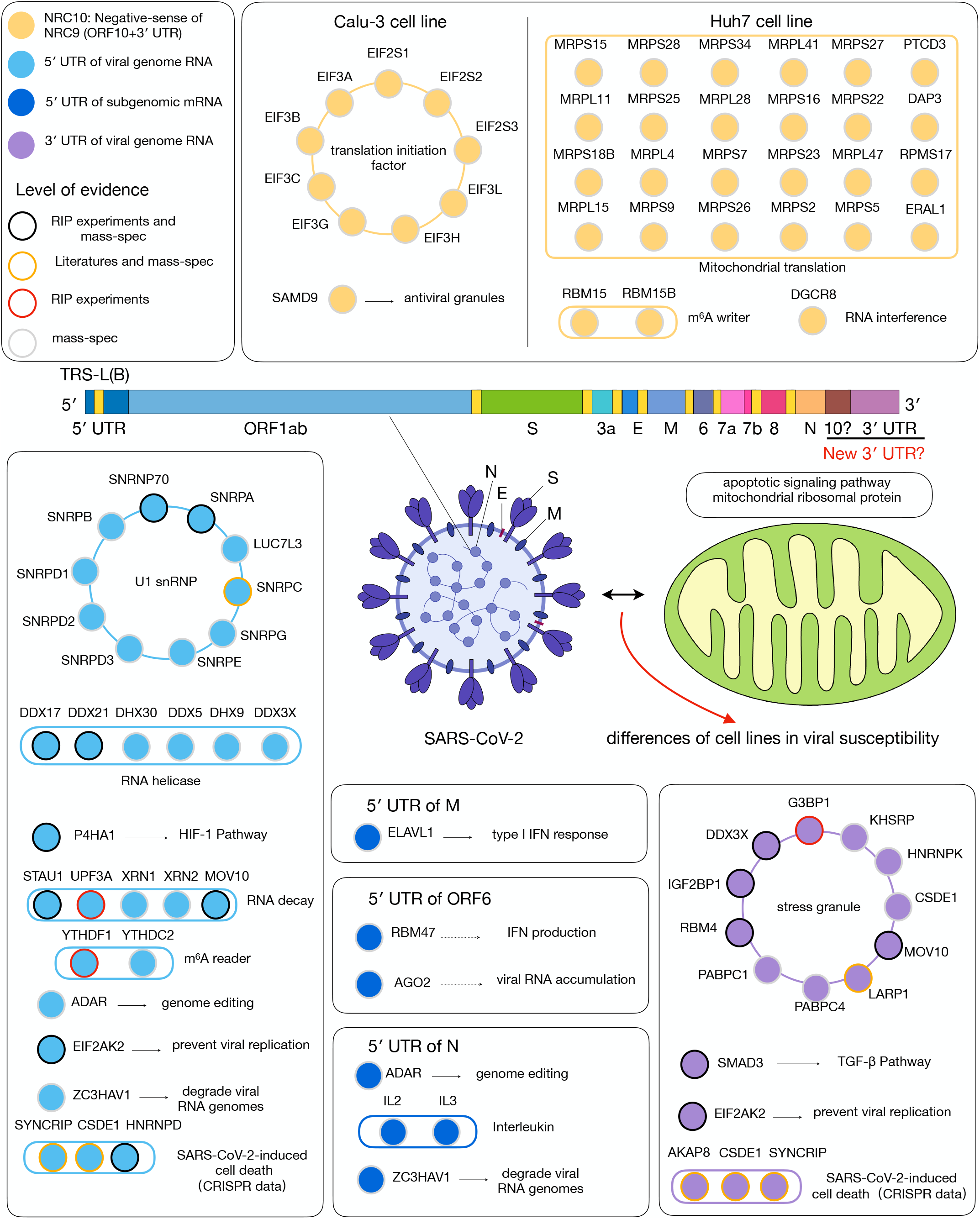
Summary of the functions of SARS-CoV-2 ncrRNA interactomes. Each node denotes a protein. Color of nodes indicates the specific group of ncrRNAs. Border color of nodes indicate the level of evidence. Solid edges represent associations based on direct evidences. Dashed edges represent associations based on indirect evidences.

In particular, the viral 5′ UTR interactome features proteins involved in the U1 snRNP and regulation of viral replication and transcription. A previous study reveals that the viral NSP16 protein can bind to the recognition domain of U1/U2 snRNAs to disrupt host mRNA splicing^78^. Thus, our findings provide a potential new mechanism through which snRNPs assist the transcription and/or replication of SARS-CoV-2. ADAR, STAU1, XRN1, XRN2, and P4HA1 were identified to bind to the 5′ UTR, suggesting host cell may regulate viral replication via genome editing, RNA decay, and HIF-1 pathways. In addition, our data uncovered the interactions between the m^6^A reader proteins (YTHDC2 and YTHDF1) and the viral 5′ UTR (Figure 2B and 3A). m^6^A is the most abundant type of methylation in mRNA, which is involved in several important biological processes, such as RNA stability, transport, and protein translation^79^. It has been reported that this modification can regulate the abundance of SARS-CoV-2^80^.

Interestingly, significant differences were observed among the 5′ UTRs of different subgenomic mRNAs. The 5′ UTR of N gene (NRC6) interacted with proteins related to the regulation of viral genome replication and response to viruses, such as ILF2, ILF3, and ZC3HAV1 (Figure S7A-E), suggesting that the 5′ UTRs of subgenomic mRNAs play important and specific roles during replication and translation.

The viral 3′ UTR had strong associations with proteins related to SGs and translation. SARS-CoV-2 modulates the proteins associated with SGs to maximize replication efficiency^78,81^. For example, N protein-mediated SG disassembly can promote the production of SARS-CoV-2^78^. Interaction between 3′ UTR and SGs may play an important role in the transcription and/or replication of SARS-CoV-2. Several proteins (CSDE1, SYNCRIP, and EIF2AK2) involved in the regulation of translation were identified to bind to 3′ UTR, 5′ UTR and 5′ UTR-L, supporting the formation of a mRNA closed-loop^61^. Importantly, we did not observe significant difference between ORF10 and 3’ UTR NRCs, providing strong evidences that ORF10 is a part of the SARS-CoV-2 3’ UTR.

We also explored the possible roles of negative-sense RNAs, especially NRC10. The interactomes of NRC10 are drastically different from the rest in both cell lines but in different ways. NRC10 can bind to diverse proteins, such as CSTF3, RBM15, and SAMD9. These findings suggest that negative-sense ncrRNAs may directly regulate replication and translation processes during infection.

Finally, we identified the cell-type-specific interactions between host proteins and ncrRNAs. Compared to Calu-3 cells, there were limited associations of SARS-CoV-2 ncrRNAs with proteins involved in autophagy-related functions in Huh7 cells. However, 28S ribosomal proteins which are found in mitochondria were only enriched in Huh7 cells. Mitochondria are important for regulating cell death and innate immune signaling, and a recent study reveals the important role of mitochondria during SARS-CoV-2 infection^11^. Our data reveal cell-type-specific interactions between viral ncrRNAs and host mitochondrial proteins, providing a potential mechanism to explain the drastic differences of viral susceptibility between the two cell lines.

In summary, our work reveals a comprehensive view of functional SARS-CoV-2 ncrRNA interactome. We believe that these findings provide a new mechanistic perspective on how the virus manipulates and interacts with host cells, and offer a new strategy for the development of effective therapeutics.

### Limitations of the study

MAMS method is an *in vitro* binding assay. The rationale was to limit the impact of viral proteins associated with infection experiments, but the expression of host proteins could change during infection. Some interactions may be missing as the proteins may not be present without prolonged infection. We also did not validate all ncrRNA-protein interaction *in vivo* and in all cell lines. In addition, we revealed several potential therapeutic targets, but future investigations of these targets are needed to verify.

## Acknowledgements

We thank all the participants for agreeing to join this study. This research was supported by grants from National Natural Science Foundation of China (32022039, 91940302, 31870810, 91640104, 31670826, 31730057, 31571447, 31371417, 91540205, 22074132, 91953103), the Fundamental Research Funds for the Central Universities and Zhejiang Province (LR19C050003), special COVID-19 program of the Sino-German Center for Research Promotion (C-0023), the outstanding youth fund of Zhejiang Province (LR20B050001), and Open Project Program of the State Key Laboratory of Proteomics (SKLPO201806).

## Author contributions

L.Z. designed the RNA constructs, purified and prepared RNA and protein samples with the assistance of C.L. under the supervision of A.R.. M.X. designed and performed biochemical and cellular experiments with the assistance of Y.L. under the supervision of X.F.. Q.L. performed the mass spectrometry experiments under the supervision of B.Y.. L.J, Q.L., L.Z., C.J., M.X., B.Y. and A.R. performed raw data analyses. L.J. performed statistical analyses and data visualizations under the supervision of C.J.. L.J., C.J., M.X., L.Z., Q.L., A.R., and B.Y. drafted the manuscript. C.J., L.J., and M.X. revised the manuscript with input from all authors.

## Conflict of interest

The authors declare that they have no conflict of interest.

## Supplementary Figure Legends

**Figure S1.**
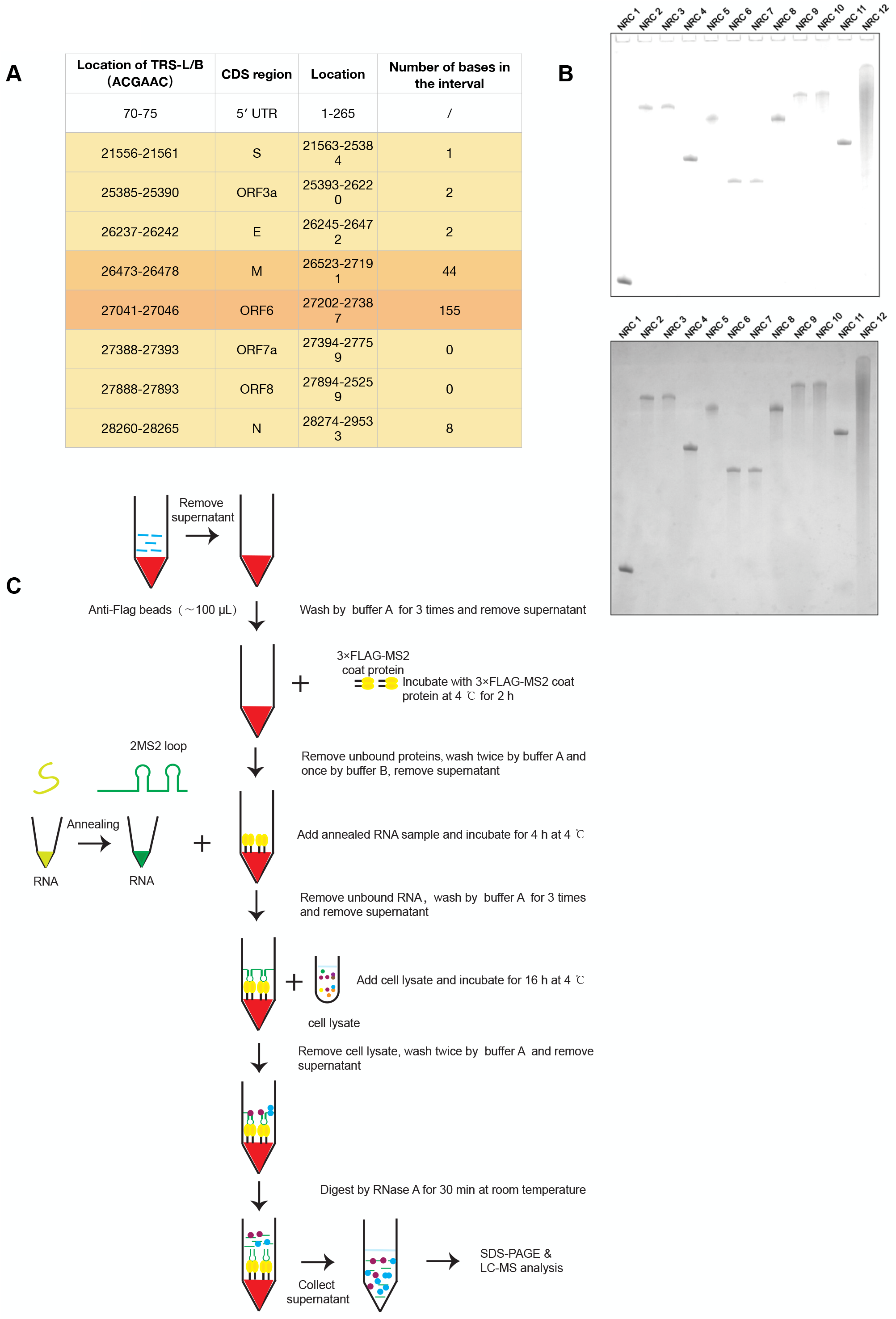
Design and validation of NRCs. (A) Location information of TRS-L/TRS-Bs. (B) Gel graphs of 12 NRCs after long (top) and short (bottom) exposures. (C) The detailed workflow for RNA pull-down assays.

**Figure S2.**
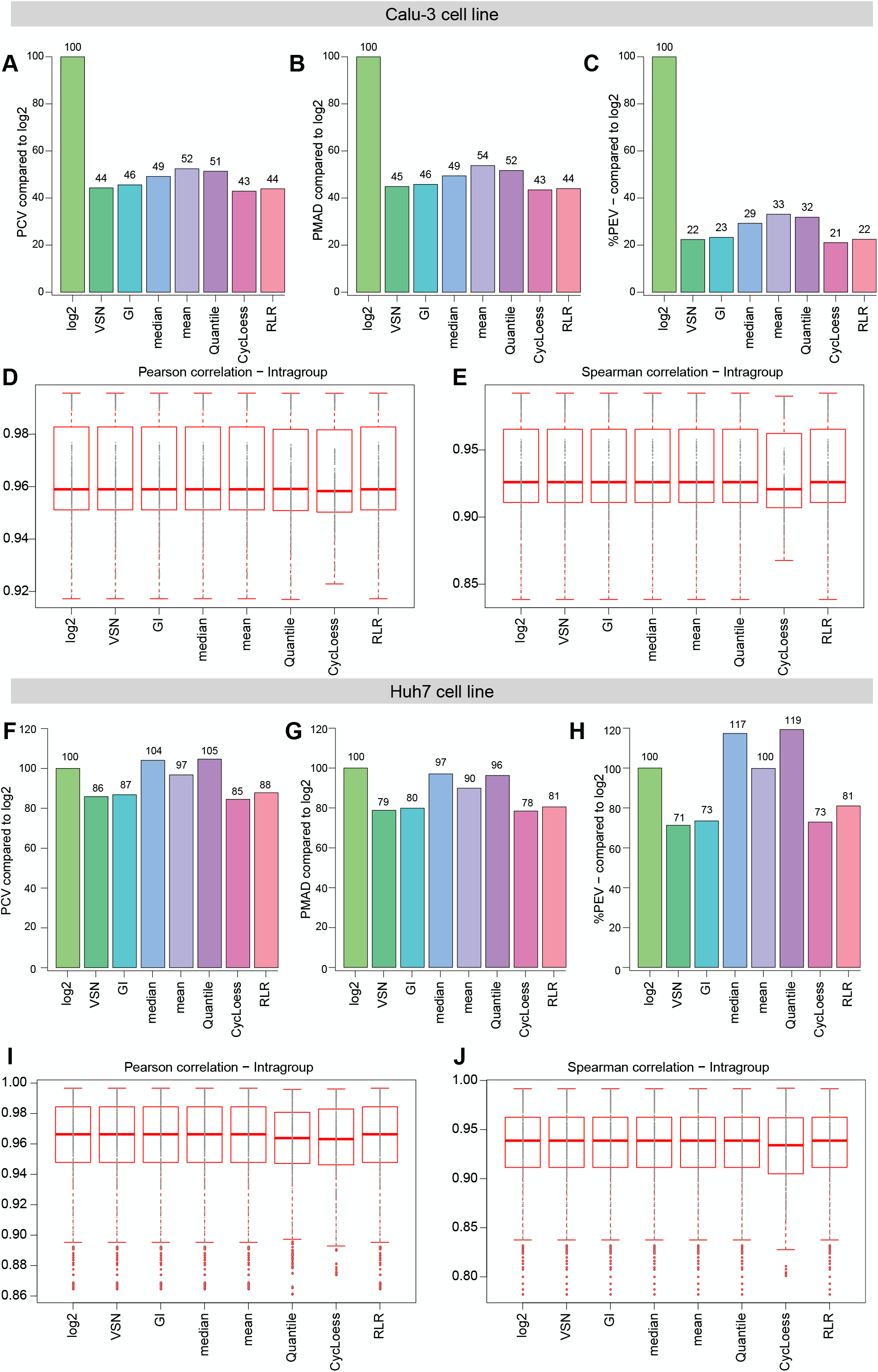
Systematic evaluation of different normalization methods. (A-C) Bar plots demonstrating the PCV (A), PMAD (B), and PEV (C) statistics of different normalization methods when compared to the log2 transformation in Calu-3 cells (The lower the better). (D-E) Box plots showing the Person correlation (D) and Spearman correlation (E) between raw data and normalized data in Calu-3 cells. (F-H) Bar plots demonstrating the PCV (F), PMAD (G), and PEV (H) statistics of different normalization methods compared to the standard log2 transformation in Huh7 cells. (I-J) Box plots showing the Person correlation (I) and Spearman correlation (J) between raw data and normalized data in Huh7 cells.

**Figure S3.**
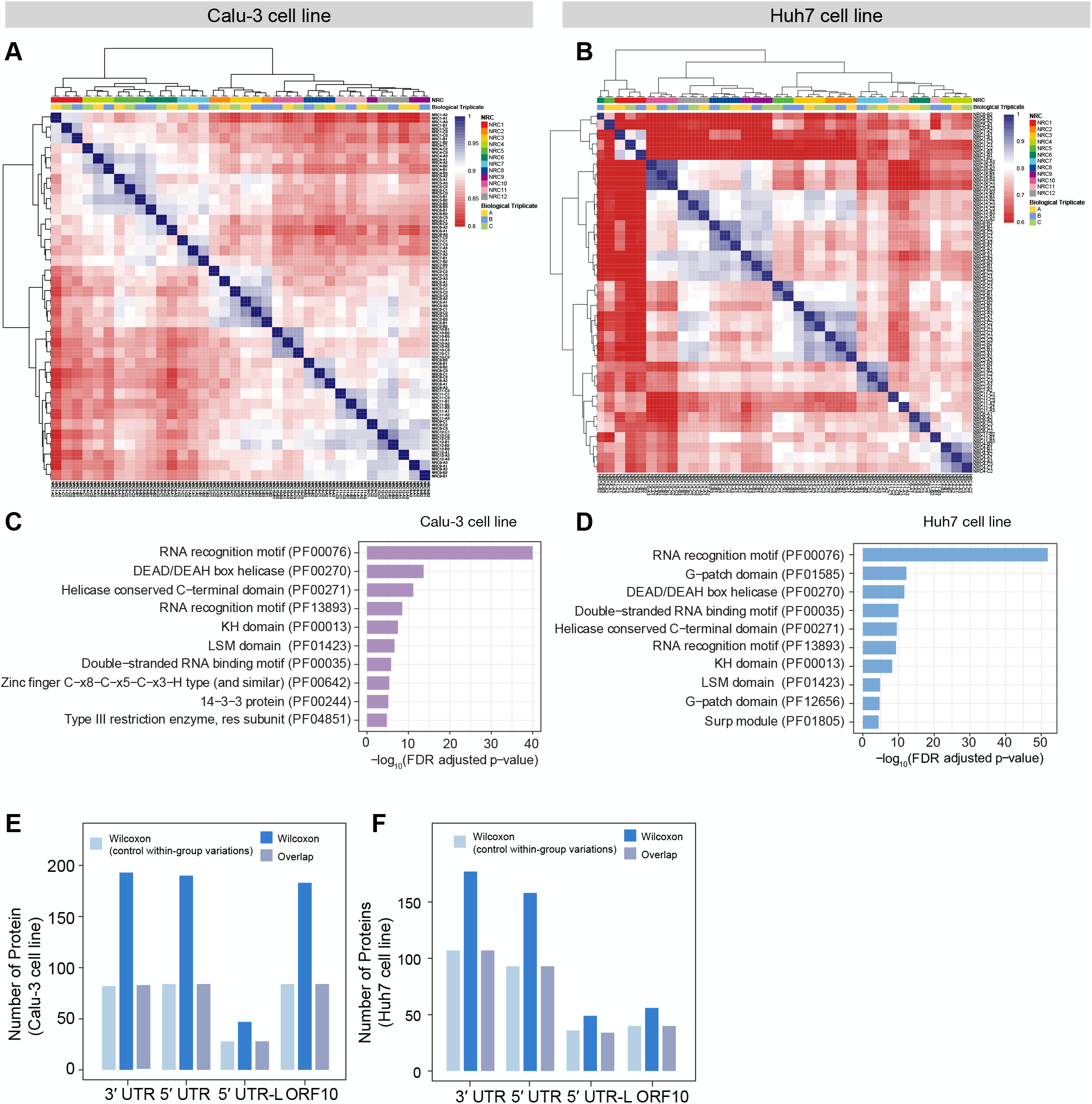
Analysis of SARS-CoV-2 ncrRNA-host protein interactome. (A-B) Heatmaps of the interactomes revealing that the biological and technical replicates of NRCs are highly correlated in both Calu-3 (A) and Huh7 (B) cells. (C-D) Protein domain enrichment analyses of host proteins enriched in SARS-CoV-2 ncrRNA interactomes in Calu-3 (C) and Huh7 (D) cells. (E-F) Bar plots showing the overlap of host proteins identified by two statistical methods in Calu-3 (E) and Huh7 (F) cells.

**Figure S4.**
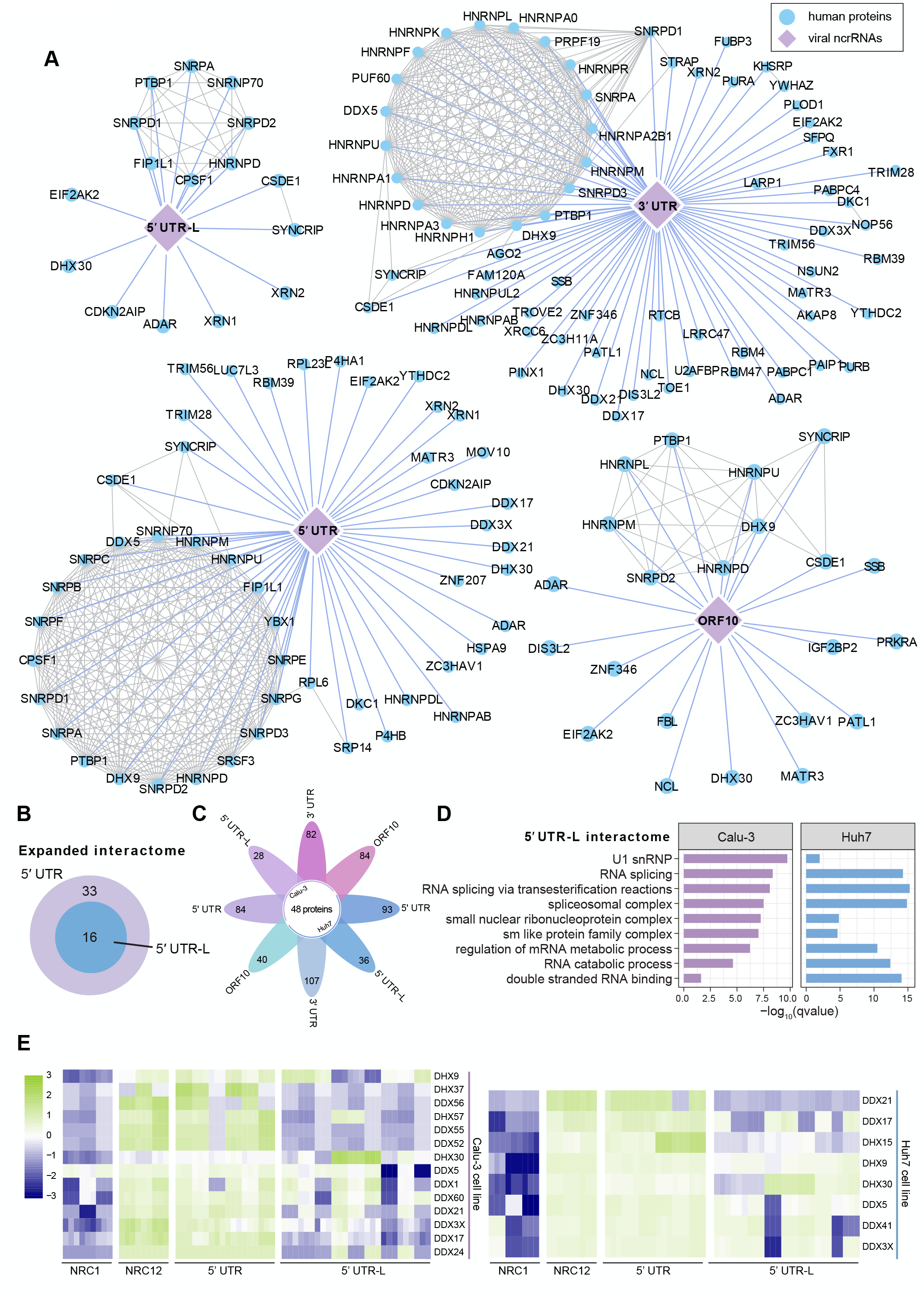
The expanded interactome of SARS-CoV-2. (A) The expanded SARS-CoV-2 ncrRNAs-host protein interactome. (B) Overlap between expanded 5′ UTR interactome and 5′ UTR-L interactome. (C) Flower plot showing the core proteins identified by two cell lines. (D) GO enrichment of 5′ UTR-L interactomes in two cell lines. (E) Heatmap showing the interactions between RNA helicases and 5**′** UTR or 5**′** UTR-L.

**Figure S5.**
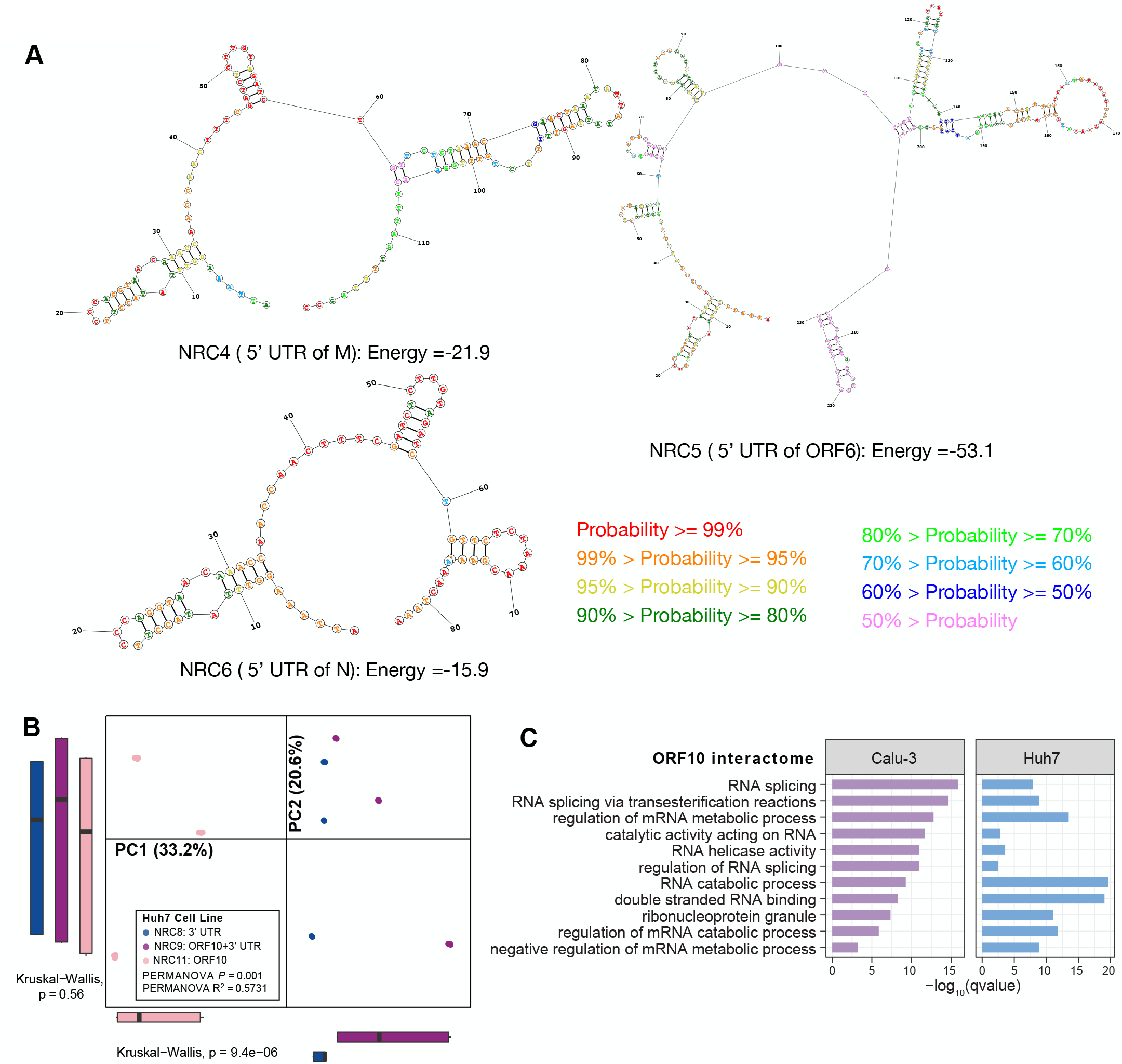
The role of SARS-CoV-2 5′UTR-L NRCs and ORF10. (A) Predicted results of secondary structures of NRC4, NRC5, and NRC6. Colors of bases (A) indicate probability of prediction. (B) PCA analysis of interactomes of the NRC8, NRC9, and NRC11 in Huh7 cells. Points represent samples. Colors represent different NRCs. (C) GO enrichment analysis of the interactome of ORF10 in two cell lines.

**Figure S6.**
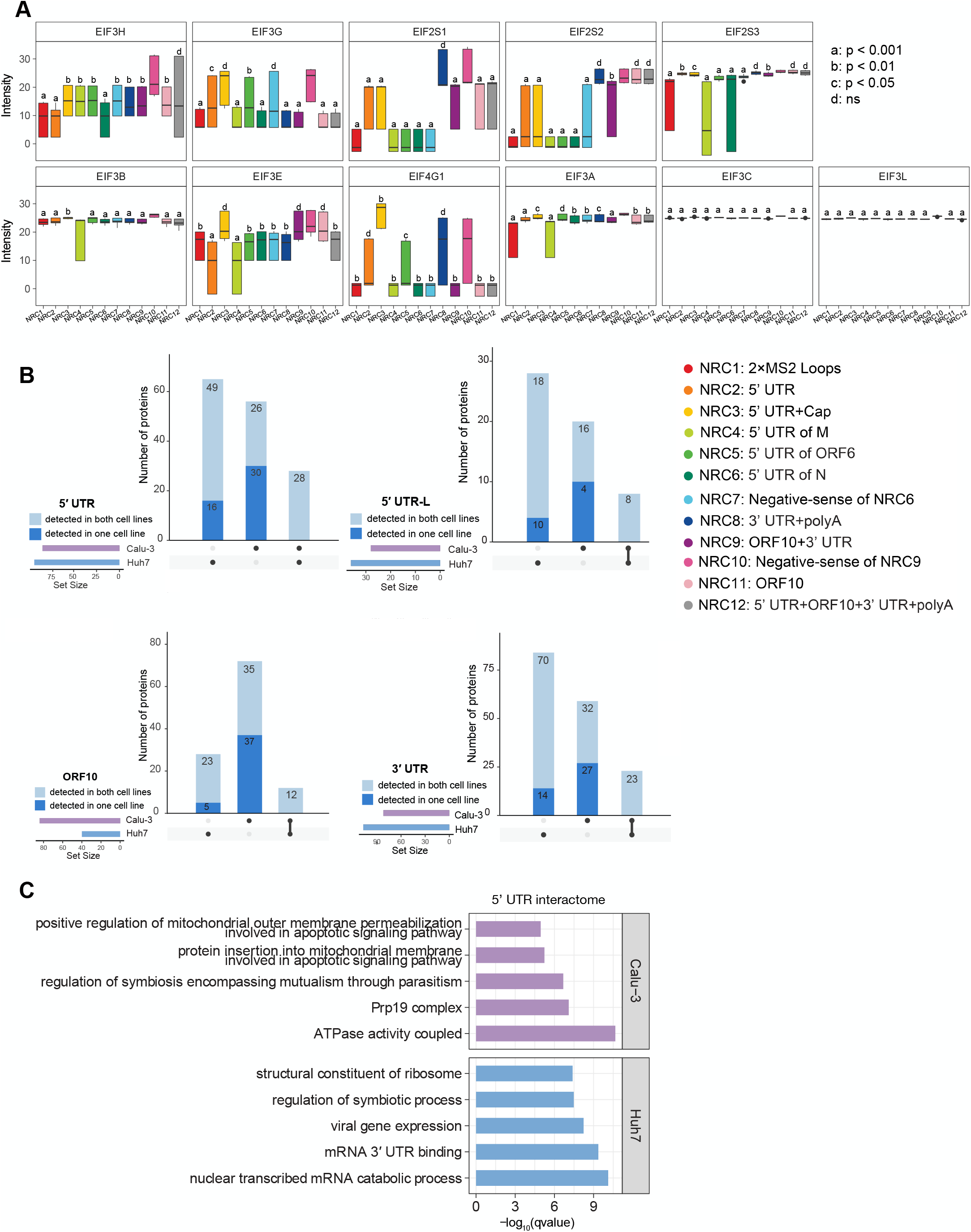
Comparison of the interactomes of Calu-3 and Huh7 cells. (A) The intensity values of translation initiation factors in Calu-3 cells. *P* values between NRC10 and other NRCs were calculated by the Wilcoxon test. a, b, c and d labels represent the *P* values: d represents ns (not significant), c represents *p* < 0.05, b represents *p* < 0.01, and a represents *p* < 0.001. (B) UpSet plots showing the cell-specific-interactions between SARS-CoV-2 ncrRNAs and host proteins. Dark blue represents the proteins detected in both cell lines. Light blue represents the proteins detected in either Calu-3 or Huh7 cells. (C) GO enrichment analysis revealing different biological functions of the 5′ UTR interactome in two cell lines.

**Figure S7.**
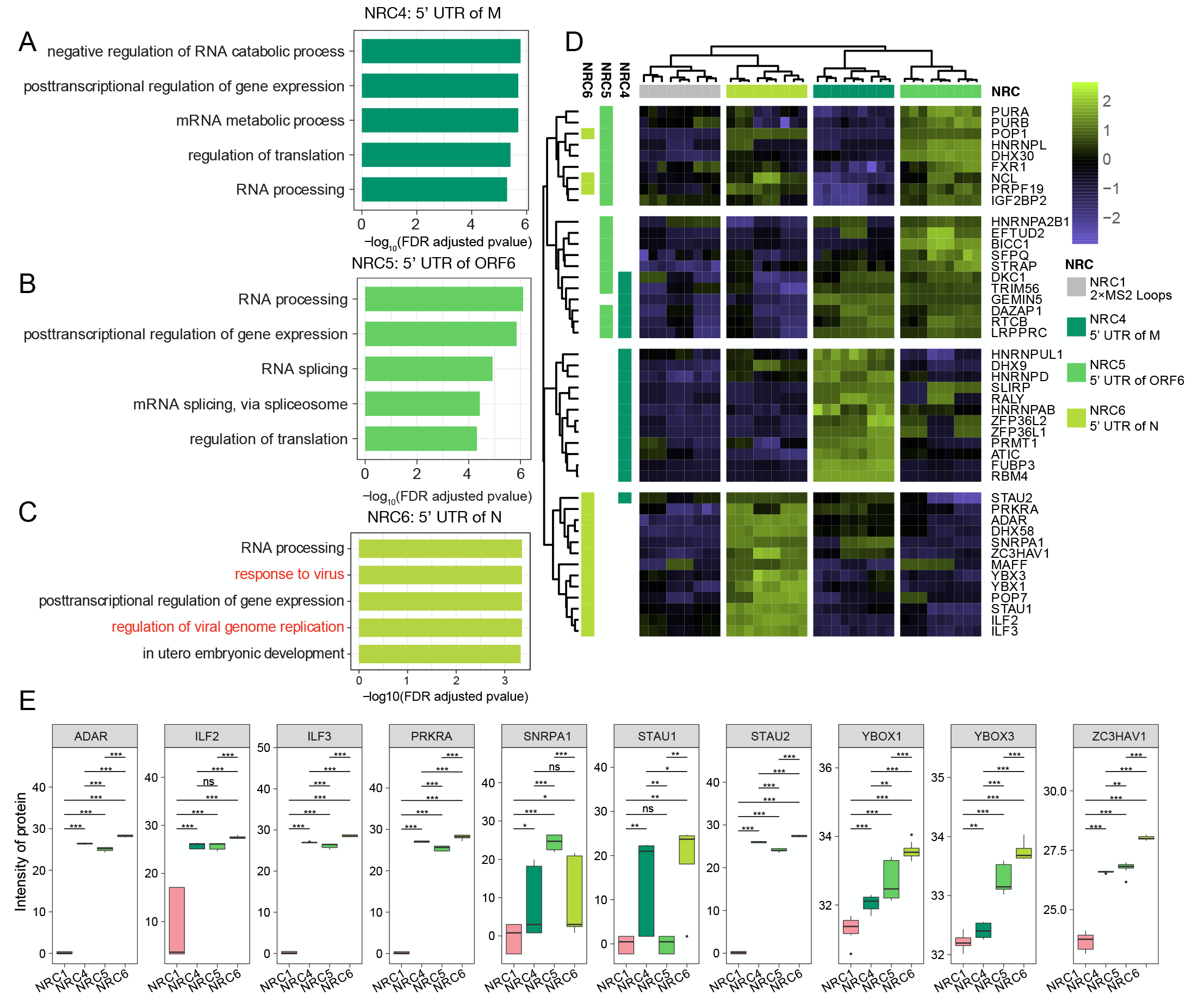
Comparison of the interactomes of 5′ UTRs of subgenomic mRNAs. (A) GO enrichment analysis of proteins bound to NRC4 specifically in Calu-3 cells. (B) GO enrichment analysis of proteins bound to NRC5 specifically in Calu-3 cells. (C) GO enrichment analysis of proteins bound to NRC4 specifically in Calu-3 cells. (D) Heatmap revealing that NRC4, NRC5, and NRC6 bind to different proteins in Calu-3 cells. Colored columns on the left denote the identity of the groups. (E) The intensity values of proteins interacted with NRC6 specifically in Huh7 cells. *P* values were calculated using the Wilcoxon test. * *p* < 0.05, ** *p* < 0.01, *** *p* < 0.001.

## Supplementary Table Legends

**Table S1. 12 NRCs for RNA pulldown assays**.

**Table S2. Mass spectrometry data for viral ncrRNA interactomes in Calu-3 and Huh7 cells:** Full data for all proteins in all datasets including proteomes of Calu3-3 and Huh7, the expanded SARS-CoV-2 ncrRNA interactome, and the core SARS-CoV-2 ncrRNA interactome.

**Table S3. Intersection of the SARS-CoV-2 ncrRNA interactome with published work. Table S4. RIP experiment data in HEK293 cells**.

## Methods

### RNA purification and preparation

The DNA sequence of each NRC (NRC1-12) was cloned into pUT7 vector bearing the T7 RNA polymerase promoter^82^ and was amplified by PCR with *pfu* DNA polymerase, which was used as the templates for transcription. The *in vitro* transcription of all the RNA samples of 12 NRCs were carried out at 37 °C by T7 RNA polymerase, followed by purification with denatured urea-PAGE (polyacrylamide gel electrophoresis) and precipitation with ethanol. The purified RNA samples were annealed at 65 °C for 5 min in the buffer containing 50 mM HEPES, pH 6.8, 50 mM NaCl and 5 mM MgCl2, followed by incubation on ice for 0.5 h∼1 h before performing RNA pull-down assays.

### *In vitro* RNA capping

NRC3 has the same sequence as NRC2, but with 5’-end 7-methylguanylate cap (cap 0 format). The capping reaction of NRC2 was performed *in vitro* with vaccinia capping enzyme (YEASEN) at 37 °C, which catalyzes the addition of 7-methylguanylate structures (cap 0) to the 5’-end of RNA. Phenol-chloroform-isoamyl alcohol extraction was performed to remove the enzyme from the reaction products. After ethanol precipitation and annealing, the capped RNA sample (NRC3) was used for RNA pull-down assays.

### Preparation of 3×FLAG-MS2 phage coat protein (FLAG-MCP)

In RNA pull-down assays, the 3×FLAG-MS2 phage coat protein mediates the binding of RNA samples to Anti-Flag beads. The protein sequence was shown as following, <u>DYKDHDGDYKDHDIDYKDDDDK</u>GGSMASNFTQFVLVDNGGTGDVTVAPSNFANGI AEWISSNSRSQAYKVTCSVRQSSAQNRKYTIKVEVPKGAWRSYLNMELTIPIFATNSDCELIVKAMQGLLKDGNPIPSAIAANSGIY, in which the 3×FLAG-tag was underlined.

For purification, one 6-his-SUMO-tag followed by one ubiquitin-like protease (ULP1) cleavage site were fused to the N-terminal of the 3×FLAG-MS2 phage coat protein. The fused protein was expressed in *Escherichia coli* BL21(DE3) Codon plus strain. The lysis of the cultured cells was carried out with French press in buffer A containing 25 mM Tris, pH 8.0, 1.0 M NaCl, 5 mM 2-mercaptoethanol, supplemented with 0.1 mM phenylmethylsulfonyl fluoride (PMSF). After centrifugation, the supernatant sample was loaded onto the first HisTrap column (GE Healthcare) and eluted by buffer B containing 25mM Tris, pH 8.0, 500 mM NaCl, 5 mM 2-mercaptoethanol and 500 mM imidazole. Then the eluted fused protein was incubated with ULP1 protease overnight for cleavage of the fused tag. The cleaved fused 6His-SUMO-tag was removed from the FLAG-MCP by reloading the sample onto the second HisTrap column (GE Healthcare). The FLAG-MCP was further purified by chromatography using a HiTrap Heparin SP column (GE Healthcare) and a HiLoad Superdex 75 16/60 column (GE Healthcare). The purified FLAG-MCP was concentrated before storage at −80 °C in the buffer containing 40 mM HEPES, pH 7.0, 50 mM KCl, 100 mM NaCl, 5 mM MgCl2 and 2 mM dithiothreitol (DTT).

### Cell lysis and digestion

Each sample was separated by SDS-PAGE gel and stained by Coomassie brilliant blue respectively. Protein gel bands were cut into 1× 1 × 1mm pieces and collected into a 1.5 mL centrifuge tube. Gel pieces were destained with 25% acetonitrile (ACN) in 50 mM ammonium bicarbonate (ABC) for 20 minutes at room temperature (RT) with shaking. The gel was dehydrated using 1 mL 100% ACN twice with shaking. Proteins were reduced with 10 Mm DTT for 60 min at 56 °C and alkylated with 55 mM iodoacetamide for 45 min. Gel pieces were washed with digestion buffer (50 mM NH4HCO3, pH 8.0) twice, dehydrated with acetonitrile, and then dry the sample with speed-vac. Gel pieces were rehydrated with trypsin solution (10 ng/◸l sequencing grade modified trypsin, 50 mM NH4HCO3, pH 8.0) and incubated overnight at 37 °C. Digested peptides were extracted from gel pieces with elution buffer 1 (50% acetonitrile, 5% formic acid), elution buffer 2 (75% acetonitrile, 0.1% formic acid), sequentially. Gel pieces were dehydrated with acetonitrile twice, and all of supernatant were combined. The peptides solution was dried with speed vac, digested peptides were resuspended with 5% formic acid and desalted with StageTip.

### RNA pull-down assays

The detailed workflow for RNA pull-down assays in this paper is shown in Figure S1C. 50 μL anti-flag beads were incubated with excess FLAG-MCP sample in the buffer A (50 mM Tris, pH7.5, 150 mM NaCl, 0.1% NP-40, and 2 mM MgCl2) at 4 °C for 2 h, followed by washing with buffer A twice to remove the unbound proteins. Then the annealed purified RNA sample was added to the FLAG-MCP bound beads and incubated at 4 °C for 4 h. The unbound RNA was removed by washing with buffer A for 3 times. In the following step, the cell lysate was added to the RNA-MCP bound beads and the incubation time was extended to 16 h at 4 °C. For each NRC, we used Huh7 and Calu-3 cell lines for experiments. After incubation, the cell lysate was removed by washing twice with buffer A. Then, RNase A was added to digest RNA molecules and release the protein components that bound with each RNA sample from the RNA-MCP bound beads. The products were analyzed with SDS-PAGE and followed with liquid chromatography-mass spectrometry. Each assay was repeated 3 times independently.

### LC-MS data acquisition

Digested peptides were analyzed on an Q Exactive HF-X mass spectrometry system (Thermo Fisher Scientific) equipped with an Easy-nLC 1200 liquid chromatography system (Thermo Fisher Scientific). Samples were injected on a C18 reverse phase column (75 ◸m × 15 cm, 1.9 ◸m C18, 5 mm tip). Mobile phase A consisted of 0.1% FA, and mobile phase B consisted of 0.1% FA/80% ACN. Peptides were analyzed with a 60 minutes linear gradient at flow rate 200 nL minutes^-1^ as the following: 0–5% B for 2 minutes, 5–35% B for 46 minutes, 35–100% B in 12 minutes. Data dependent analysis (DDA) was performed by acquiring a full scan over a m/z range of 350-1500 in the Orbitrap at R = 60,000 (m/z = 200), NCE = 27, with a normalized AGC target of 3 × 10^6^, an isolation width of 0.8 m/z. The AGC targets and maximum ion injection time for the MS2 scans were 3 × 10^5^ and 60 ms, respectively. Precursors of the + 1, + 8 or above, or unassigned charge states were rejected; exclusion of isotopes was disabled; dynamic exclusion was set to 45 s. Mass spectrometry data were searched by MaxQuant (Version 1.6.10.43).

### Quality control and assessment of LC-MS data

Raw mass spectrometric data files were analyzed by MaxQuant software. All data were searched against the SwissProt Human protein sequences. Peptide and protein identification as well as label-free quantitation were performed; false-discovery rate (FDR) was set to 1%; fixed modification was carbamidomethyl; main search peptide tolerance was 10 ppm.

MaxQuant outputs were used for downstream analysis. For each biological replicate, proteins that meet any of the following criteria are filtered out: 1. flagged as potential contaminants; 2. flagged as reverse sequences; 3. only identified by site; 4. quantified by a single razor or unique peptide; and 5. only detected once in all samples. Missing data imputation was performed for proteins with missing values in one technical replicate, while present in the other two replicates. Imputation was performed using mean value of the other two technical replicates. Finally, 638 and 449 proteins were detected in Calu-3 and Huh7 cells, respectively. For data normalization, NormalyzerDE (version 1.5.4)^21^ was applied to select the best normalization method. Based the results of pooled estimate of variance (PEV), coefficient of variation (CV), median absolute deviation (MAD), and correlation analyses, we chose the VSN as the normalization method (Figure S2). To remove the batch effects and correct data, we performed the ComBat method using the R package ‘sva’ (version 3.38.0)^22^. Principle component analysis (PCA) was applied to examine the batch effects and the separation of interactomes across samples and using the R package ‘ade4’ (version 1.7-16)^83^. To determine the statistical significances in dissimilarity matrices across samples, the adonis test using the adonis function (999 permutations) in the R package ‘ade4‘ (version 1.7-16)^83^ was performed. To compare the statistical significances in dissimilarity of PC1 and PC2, the Kruskal-Wallis test was performed. Pairwise Spearman’ s correlation coefficients were calculated for all samples.

### Identification of viral ncrRNA-binding proteins

To identify the viral ncrRNA-binding proteins, two statistical methods were used: 1. The Wilcoxon test was performed to compare the interactomes of NRC1 and other NRCs. *P* values were adjusted using the FDR correction by the R package ‘fdrtool’ (version 1.2.16)^84^. Statistical significance was set as adjusted *P* value < 0.05; and 2. The Wilcoxon test was applied to identify the proteins enriched in 5′ UTR, 5′ UTR-L, 3′ UTR and ORF10 groups. To control within-group variations, between-group variation/ within-group variation was set as > 2. *P* values were adjusted using the FDR correction by the R package ‘fdrtool’ (version 1.2.16)^84^. Statistical significance was set as adjusted *P* value < 0.05.

### RIP experiment

HEK 293T cells were transfected with plasmids co-expressing individual RNA binding protein, MS2, 5‘UTR and 3’ UTR. Cells were harvested and resuspended in lysis buffer (50 mM Tris-HCl [pH7.5], 150 mM NaCl, 1% NP-40, 2 mM MgCl2). Cell lysate was incubated with anti-FLAG M2 Magnetic Beads (Sigma, #M8823) at 4 °C for 4 h. After 5 times washes by lysis buffer, beads were collected. Recovered RNA is isolated by Trizol. Purified RNAs were detected by quantitative real time PCR.

### Enrichment analyses

Sets of proteins bound to viral ncrRNAs were tested for enrichment of pathways using the Reactome web interface (https://reactome.org/)^23^. Statistical significance was set as FDR adjusted *P* value < 0.05. Gene Ontology (GO) enrichment analysis was carried out using the R package ‘clusterProfiler’ (version 3.18.0)^85^. The gene ontology terms were obtained from the c5 category of Molecular Signature Database using R package ‘msigdb’ (version 7.2)^86^. Statistical significance was set as qvalue < 0.05. Protein domain enrichment analysis was carried out using the STRING web interface (https://string-db.org/)^87^. Statistical significance was set as FDR adjusted *P* value < 0.05.

### Protein-Protein interaction network analysis

Protein-protein interactions network was performed using the STRING web interface (https://string-db.org/)^87^. Minimum required interaction score was set as the highest confidence (0.900). Cytoscape software (version 3.8.2)^29^ was applied to visualize the network.

### Prediction of RNA secondary structure

RNAstructure (http://rna.urmc.rochester.edu/RNAstructureWeb/)^47^ was applied to predict the secondary structure of RNA using the default parameters.

## Data visualization

Most of the data visualization were performed in R using the packages “ggplot2” (version 3.3.3)^88^, “pheatmap” (version 1.0.12)^89^, “ComplexHeatmap” (version 2.6.2)^90^ and “UpsetR” (version 1.4.0)^91^. E Venn website (http://www.ehbio.com/test/venn/#/) was applied to create Venn diagram and flower plot.

## Notes

### Competing Interest Statement

The authors have declared no competing interest.

## Reference

1. Guan WJ, Ni ZY, Hu Y, et al. Clinical Characteristics of Coronavirus Disease 2019 in China. N Engl J Med. 2020;382(18):1708–1720.

2. Dong E, Du H, Gardner L. An interactive web-based dashboard to track COVID-19 in real time. Lancet Infect Dis. 2020;20(5):533–534.

3. da Silva SJR, Alves da Silva CT, Mendes RPG, Pena L. Role of nonstructural proteins in the pathogenesis of SARS-CoV-2. J Med Virol. 2020;92(9):1427–1429.

4. Lu R, Zhao X, Li J, et al. Genomic characterisation and epidemiology of 2019 novel coronavirus: implications for virus origins and receptor binding. Lancet. 2020;395(10224):565–574.

5. Zhang YZ, Holmes EC. A Genomic Perspective on the Origin and Emergence of SARS-CoV-2. Cell. 2020;181(2):223–227.

6. V’Kovski P, Kratzel A, Steiner S, Stalder H, Thiel V. Coronavirus biology and replication: implications for SARS-CoV-2. Nat Rev Microbiol. 2021;19(3):155–170.

7. Sola I, Almazan F, Zuniga S, Enjuanes L. Continuous and Discontinuous RNA Synthesis in Coronaviruses. Annu Rev Virol. 2015;2(1):265–288.

8. Sawicki SG, Sawicki DL. Coronaviruses use discontinuous extension for synthesis of subgenome-length negative strands. Adv Exp Med Biol. 1995;380:499–506.

9. Lai MM, Stohlman SA. Comparative analysis of RNA genomes of mouse hepatitis viruses. J Virol. 1981;38(2):661–670.

10. Schmidt N, Lareau CA, Keshishian H, et al. The SARS-CoV-2 RNA-protein interactome in infected human cells. Nat Microbiol. 2021;6(3):339–353.

11. Flynn RA, Belk JA, Qi Y, et al. Discovery and functional interrogation of SARS-CoV-2 RNA-host protein interactions. Cell. 2021;184(9):2394–2411 e2316.

12. Kamel W, Noerenberg M, Cerikan B, et al. Global analysis of protein-RNA interactions in SARS-CoV-2 infected cells reveals key regulators of infection. bioRxiv. 2020:2020.2011.2025.398008.

13. Lee S, Lee Y-s, Choi Y, et al. The SARS-CoV-2 RNA interactome. bioRxiv. 2020:2020.2011.2002.364497.

14. Chan AP, Choi Y, Schork NJ. Conserved Genomic Terminals of SARS-CoV-2 as Coevolving Functional Elements and Potential Therapeutic Targets. mSphere. 2020;5(6).

15. Thi Nhu Thao T, Labroussaa F, Ebert N, et al. Rapid reconstruction of SARS-CoV-2 using a synthetic genomics platform. Nature. 2020;582(7813):561–565.

16. Kim D, Lee JY, Yang JS, Kim JW, Kim VN, Chang H. The Architecture of SARS-CoV-2 Transcriptome. Cell. 2020;181(4):914–921 e910.

17. Wu F, Zhao S, Yu B, et al. A new coronavirus associated with human respiratory disease in China. Nature. 2020;579(7798):265–269.

18. Pancer K, Milewska A, Owczarek K, et al. The SARS-CoV-2 ORF10 is not essential in vitro or in vivo in humans. PLoS Pathog. 2020;16(12):e1008959.

19. Davidson AD, Williamson MK, Lewis S, et al. Characterisation of the transcriptome and proteome of SARS-CoV-2 reveals a cell passage induced in-frame deletion of the furin-like cleavage site from the spike glycoprotein. Genome Med. 2020;12(1):68.

20. Yao H, Lu X, Chen Q, et al. Patient-derived SARS-CoV-2 mutations impact viral replication dynamics and infectivity in vitro and with clinical implications in vivo. Cell Discov. 2020;6(1):76.

21. Willforss J, Chawade A, Levander F. NormalyzerDE: Online Tool for Improved Normalization of Omics Expression Data and High-Sensitivity Differential Expression Analysis. J Proteome Res. 2019;18(2):732–740.

22. Leek JT, Johnson WE, Parker HS, Jaffe AE, Storey JD. The sva package for removing batch effects and other unwanted variation in high-throughput experiments. Bioinformatics. 2012;28(6):882–883.

23. Jassal B, Matthews L, Viteri G, et al. The reactome pathway knowledgebase. Nucleic Acids Res. 2020;48(D1):D498–D503.

24. Hadjadj J, Yatim N, Barnabei L, et al. Impaired type I interferon activity and inflammatory responses in severe COVID-19 patients. Science. 2020;369(6504):718–724.

25. Spiegel M, Pichlmair A, Muhlberger E, Haller O, Weber F. The antiviral effect of interferon-beta against SARS-coronavirus is not mediated by MxA protein. J Clin Virol. 2004;30(3):211–213.

26. Sallard E, Lescure FX, Yazdanpanah Y, Mentre F, Peiffer-Smadja N. Type 1 interferons as a potential treatment against COVID-19. Antiviral Res. 2020;178:104791.

27. Hensley LE, Fritz LE, Jahrling PB, Karp CL, Huggins JW, Geisbert TW. Interferon-beta 1a and SARS coronavirus replication. Emerg Infect Dis. 2004;10(2):317–319.

28. Makjaroen J, Somparn P, Hodge K, Poomipak W, Hirankarn N, Pisitkun T. Comprehensive Proteomics Identification of IFN-lambda3-regulated Antiviral Proteins in HBV-transfected Cells. Mol Cell Proteomics. 2018;17(11):2197–2215.

29. Shannon P, Markiel A, Ozier O, et al. Cytoscape: a software environment for integrated models of biomolecular interaction networks. Genome Res. 2003;13(11):2498–2504.

30. Heinrichs V, Bach M, Winkelmann G, Luhrmann R. U1-specific protein C needed for efficient complex formation of U1 snRNP with a 5’ splice site. Science. 1990;247(4938):69–72.

31. Hu X, Harvey SE, Zheng R, et al. The RNA-binding protein AKAP8 suppresses tumor metastasis by antagonizing EMT-associated alternative splicing. Nat Commun. 2020;11(1):486.

32. Parada CA, Roeder RG. A novel RNA polymerase II-containing complex potentiates Tat-enhanced HIV-1 transcription. EMBO J. 1999;18(13):3688–3701.

33. Kerns JA, Emerman M, Malik HS. Positive selection and increased antiviral activity associated with the PARP-containing isoform of human zinc-finger antiviral protein. PLoS Genet. 2008;4(1):e21.

34. Hayakawa S, Shiratori S, Yamato H, et al. ZAPS is a potent stimulator of signaling mediated by the RNA helicase RIG-I during antiviral responses. Nat Immunol. 2011;12(1):37–44.

35. Zhu Y, Chen G, Lv F, et al. Zinc-finger antiviral protein inhibits HIV-1 infection by selectively targeting multiply spliced viral mRNAs for degradation. Proc Natl Acad Sci U S A. 2011;108(38):15834–15839.

36. Cuevas RA, Ghosh A, Wallerath C, Hornung V, Coyne CB, Sarkar SN. MOV10 Provides Antiviral Activity against RNA Viruses by Enhancing RIG-I-MAVS-Independent IFN Induction. J Immunol. 2016;196(9):3877–3886.

37. Nchioua R, Kmiec D, Muller JA, et al. SARS-CoV-2 Is Restricted by Zinc Finger Antiviral Protein despite Preadaptation to the Low-CpG Environment in Humans. mBio. 2020;11(5).

38. Baserga S, Steitz J. The RNA world. In: Gesteland RFAJF, editor: Cold Spring Harbor Laboratory Press; 1993.

39. Di Giorgio S, Martignano F, Torcia MG, Mattiuz G, Conticello SG. Evidence for host-dependent RNA editing in the transcriptome of SARS-CoV-2. Sci Adv. 2020;6(25):eabb5813.

40. Brogna S, Wen J. Nonsense-mediated mRNA decay (NMD) mechanisms. Nat Struct Mol Biol. 2009;16(2):107–113.

41. Park E, Maquat LE. Staufen-mediated mRNA decay. Wiley Interdiscip Rev RNA. 2013;4(4):423–435.

42. Nagarajan VK, Jones CI, Newbury SF, Green PJ. XRN 5’-->3’ exoribonucleases: structure, mechanisms and functions. Biochim Biophys Acta. 2013;1829(6-7):590–603.

43. Serebrovska ZO, Chong EY, Serebrovska TV, Tumanovska LV, Xi L. Hypoxia, HIF-1alpha, and COVID-19: from pathogenic factors to potential therapeutic targets. Acta Pharmacol Sin. 2020;41(12):1539–1546.

44. Xiong G, Stewart RL, Chen J, et al. Collagen prolyl 4-hydroxylase 1 is essential for HIF-1alpha stabilization and TNBC chemoresistance. Nat Commun. 2018;9(1):4456.

45. Herdy B, Karonitsch T, Vladimer GI, et al. The RNA-binding protein HuR/ELAVL1 regulates IFN-beta mRNA abundance and the type I IFN response. Eur J Immunol. 2015;45(5):1500–1511.

46. Lu LF, Zhang C, Zhou XY, et al. Zebrafish RBM47 Promotes Lysosome-Dependent Degradation of MAVS to Inhibit IFN Induction. J Immunol. 2020;205(7):1819–1829.

47. Bellaousov S, Reuter JS, Seetin MG, Mathews DH. RNAstructure: Web servers for RNA secondary structure prediction and analysis. Nucleic Acids Res. 2013;41(Web Server issue):W471–474.

48. Miao Z, Tidu A, Eriani G, Martin F. Secondary structure of the SARS-CoV-2 5’-UTR. RNA Biol. 2021;18(4):447–456.

49. Guo AX, Cui JJ, Wang LY, Yin JY. The role of CSDE1 in translational reprogramming and human diseases. Cell Commun Signal. 2020;18(1):14.

50. Boussadia O, Niepmann M, Creancier L, Prats AC, Dautry F, Jacquemin-Sablon H. Unr is required in vivo for efficient initiation of translation from the internal ribosome entry sites of both rhinovirus and poliovirus. J Virol. 2003;77(6):3353–3359.

51. Lee KM, Chen CJ, Shih SR. Regulation Mechanisms of Viral IRES-Driven Translation. Trends Microbiol. 2017;25(7):546–561.

52. Phillips SL, Soderblom EJ, Bradrick SS, Garcia-Blanco MA. Identification of Proteins Bound to Dengue Viral RNA In Vivo Reveals New Host Proteins Important for Virus Replication. mBio. 2016;7(1):e01865–01815.

53. Choi KS, Mizutani A, Lai MM. SYNCRIP, a member of the heterogeneous nuclear ribonucleoprotein family, is involved in mouse hepatitis virus RNA synthesis. J Virol. 2004;78(23):13153–13162.

54. Cassady KA, Gross M. The herpes simplex virus type 1 U(S)11 protein interacts with protein kinase R in infected cells and requires a 30-amino-acid sequence adjacent to a kinase substrate domain. J Virol. 2002;76(5):2029–2035.

55. Kang JI, Kwon SN, Park SH, et al. PKR protein kinase is activated by hepatitis C virus and inhibits viral replication through translational control. Virus Res. 2009;142(1-2):51–56.

56. Lin SS, Lee DC, Law AH, Fang JW, Chua DT, Lau AS. A role for protein kinase PKR in the mediation of Epstein-Barr virus latent membrane protein-1-induced IL-6 and IL-10 expression. Cytokine. 2010;50(2):210–219.

57. Chang JH, Kato N, Muroyama R, et al. Double-stranded RNA-activated protein kinase inhibits hepatitis C virus replication but may be not essential in interferon treatment. Liver Int. 2010;30(2):311–318.

58. Park IH, Baek KW, Cho EY, Ahn BY. PKR-dependent mechanisms of interferon-alpha for inhibiting hepatitis B virus replication. Mol Cells. 2011;32(2):167–172.

59. Okonski KM, Samuel CE. Stress granule formation induced by measles virus is protein kinase PKR dependent and impaired by RNA adenosine deaminase ADAR1. J Virol. 2013;87(2):756–766.

60. Zhang L, Alter HJ, Wang H, et al. The modulation of hepatitis C virus 1a replication by PKR is dependent on NF-kB mediated interferon beta response in Huh7.5.1 cells. Virology. 2013;438(1):28–36.

61. Tomek W, Wollenhaupt K. The “closed loop model” in controlling mRNA translation during development. Anim Reprod Sci. 2012;134(1-2):2–8.

62. Mukherjee J, Hermesh O, Eliscovich C, et al. beta-Actin mRNA interactome mapping by proximity biotinylation. Proc Natl Acad Sci U S A. 2019;116(26):12863–12872.

63. Gao B, Gong X, Fang S, et al. Inhibition of anti-viral stress granule formation by coronavirus endoribonuclease nsp15 ensures efficient virus replication. PLoS Pathog. 2021;17(2):e1008690.

64. Lu S, Ye Q, Singh D, et al. The SARS-CoV-2 nucleocapsid phosphoprotein forms mutually exclusive condensates with RNA and the membrane-associated M protein. Nat Commun. 2021;12(1):502.

65. Luo L, Li Z, Zhao T, et al. SARS-CoV-2 nucleocapsid protein phase separates with G3BPs to disassemble stress granules and facilitate viral production. Sci Bull (Beijing). 2021.

66. Yang P, Mathieu C, Kolaitis RM, et al. G3BP1 Is a Tunable Switch that Triggers Phase Separation to Assemble Stress Granules. Cell. 2020;181(2):325–345 e328.

67. Zi Z, Chapnick DA, Liu X. Dynamics of TGF-beta/Smad signaling. FEBS Lett. 2012;586(14):1921–1928.

68. Zhao X, Nicholls JM, Chen YG. Severe acute respiratory syndrome-associated coronavirus nucleocapsid protein interacts with Smad3 and modulates transforming growth factor-beta signaling. J Biol Chem. 2008;283(6):3272–3280.

69. Lunde BM, Moore C, Varani G. RNA-binding proteins: modular design for efficient function. Nat Rev Mol Cell Biol. 2007;8(6):479–490.

70. Glisovic T, Bachorik JL, Yong J, Dreyfuss G. RNA-binding proteins and post-transcriptional gene regulation. FEBS Lett. 2008;582(14):1977–1986.

71. Liu J, Xu YP, Li K, et al. The m(6)A methylome of SARS-CoV-2 in host cells. Cell Res. 2021;31(4):404–414.

72. Liu J, McFadden G. SAMD9 is an innate antiviral host factor with stress response properties that can be antagonized by poxviruses. J Virol. 2015;89(3):1925–1931.

73. Nounamo B, Li Y, O’Byrne P, Kearney AM, Khan A, Liu J. An interaction domain in human SAMD9 is essential for myxoma virus host-range determinant M062 antagonism of host anti-viral function. Virology. 2017;503:94–102.

74. Harrison AG, Lin T, Wang P. Mechanisms of SARS-CoV-2 Transmission and Pathogenesis. Trends Immunol. 2020;41(12):1100–1115.

75. Marjot T, Webb GJ, Barritt ASt, et al. COVID-19 and liver disease: mechanistic and clinical perspectives. Nat Rev Gastroenterol Hepatol. 2021;18(5):348–364.

76. Van den Broeke C, Jacob T, Favoreel HW. Rho’ing in and out of cells: viral interactions with Rho GTPase signaling. Small GTPases. 2014;5:e28318.

77. Hou L, Dong J, Zhu S, et al. Seneca valley virus activates autophagy through the PERK and ATF6 UPR pathways. Virology. 2019;537:254–263.

78. Banerjee AK, Blanco MR, Bruce EA, et al. SARS-CoV-2 Disrupts Splicing, Translation, and Protein Trafficking to Suppress Host Defenses. Cell. 2020;183(5):1325–1339 e1321.

79. Jiang X, Liu B, Nie Z, et al. The role of m6A modification in the biological functions and diseases. Signal Transduct Target Ther. 2021;6(1):74.

80. Zhang T, Yang Y, Xiang Z, et al. N(6)-methyladenosine regulates RNA abundance of SARS-CoV-2. Cell Discov. 2021;7(1):7.

81. Savastano A, Ibanez de Opakua A, Rankovic M, Zweckstetter M. Nucleocapsid protein of SARS-CoV-2 phase separates into RNA-rich polymerase-containing condensates. Nat Commun. 2020;11(1):6041.

82. Pikovskaya O, Serganov AA, Polonskaia A, Serganov A, Patel DJ. Preparation and crystallization of riboswitch-ligand complexes. Methods Mol Biol. 2009;540:115–128.

83. Dray S, Dufour A-B. The ade4 package: implementing the duality diagram for ecologists. Journal of statistical software. 2007;22(4):1–20.

84. Strimmer K. fdrtool: a versatile R package for estimating local and tail area-based false discovery rates. Bioinformatics. 2008;24(12):1461–1462.

85. Yu G, Wang L-G, Han Y, He Q-Y. clusterProfiler: an R package for comparing biological themes among gene clusters. Omics: a journal of integrative biology. 2012;16(5):284–287.

86. Liberzon A, Birger C, Thorvaldsdottir H, Ghandi M, Mesirov JP, Tamayo P. The Molecular Signatures Database (MSigDB) hallmark gene set collection. Cell Syst. 2015;1(6):417–425.

87. Szklarczyk D, Gable AL, Lyon D, et al. STRING v11: protein-protein association networks with increased coverage, supporting functional discovery in genome-wide experimental datasets. Nucleic Acids Res. 2019;47(D1):D607–D613.

88. Wickham H. ggplot2. Wiley Interdisciplinary Reviews: Computational Statistics. 2011;3(2):180–185.

89. Kolde R, Kolde MR. Package ‘pheatmap’. R package. 2015;1(7):790.

90. Gu Z, Eils R, Schlesner M. Complex heatmaps reveal patterns and correlations in multidimensional genomic data. Bioinformatics. 2016;32(18):2847–2849.

91. Conway JR, Lex A, Gehlenborg N. UpSetR: an R package for the visualization of intersecting sets and their properties. Bioinformatics. 2017;33(18):2938–2940.

